# Hydraulic modelling reveals untreated sewage, not pharmaceutical waste, drives antimicrobial resistance in a small river running through a big city

**DOI:** 10.1101/2024.12.21.629897

**Authors:** Vikas Sonkar, Arun Kashyap, Rebeca Pallares-Vega, Sai Sugitha Sasidharan, Ankit Modi, Cansu Uluseker, Sangeetha Chandrakalabai Jambu, Pranab Kumar Mohapatra, Joshua Larsen, David W Graham, Shashidhar Thatikonda, Jan-Ulrich Kreft, the AMRflows consortium

## Abstract

Quantifying sources of antimicrobial resistance (AMR) in rivers receiving various waste streams is essential for targeting mitigation strategies yet rarely performed. This study combined field monitoring with hydraulic modelling and mass balance calculations to attribute sources of AMR in the Musi River running through Hyderabad, India, a city renowned for pharmaceutical manufacturing. We quantified antibiotic resistance genes (ARGs), resistant bacteria (ARBs), and physicochemical parameters in water and sediment samples in the dry and wet season. Absolute ARG and ARB abundances spiked in the city, declining again downstream. Changes were more gradual in the wet season. Pollution levels were significantly different between upstream, city and downstream stretches and seasons. Hydraulic modelling revealed that 60-80% or 20-40% of the river water in the city derived from untreated sewage during the dry or wet seasons, respectively. This established municipal waste, not pharmaceutical sources, as the dominant driver of AMR in the Musi. Linear discriminant analysis identified dissolved oxygen and total nitrogen as reliable proxies for distinguishing AMR-polluted from less-polluted sites. The Musi, with insufficient wastewater treatment and limited dilution of point-source loadings, is typical for many urban rivers in resource-limited countries highlighting the urgent need for improved wastewater management to reduce AMR exposures.

## 1. Introduction

Antimicrobial resistance (AMR) is among the greatest health threats of the 21^st^ century, with an estimated 1.14 (95% UI 1.00–1.28) million deaths attributable to bacterial AMR in 2021, projected to rise to 1.91 (1.56–2.26) million deaths by 2050^1,2^. Although the primary human health risk stems from increasing resistance in human pathogens, their resistance is primarily conferred by antimicrobial resistance genes (ARGs) that can be acquired through horizontal gene transfer (HGT) from any antimicrobial resistant bacteria (ARBs) *via* mobile genetic elements (MGEs)^3^. Consequently, ARGs, ARBs and MGEs are often used as surrogates for monitoring AMR spread across One Health sectors.

AMR prevalence is increasing disproportionately in Low- and Middle-Income Countries (LMICs) due to inadequate waste management and less regulated antimicrobial use, resulting in AMR-laden wastes released into the environment, fuelling AMR transmission and spread *via* polluted water across One Health sectors^4–7^. In parallel, many LMICs are experiencing accelerated urbanisation that strains civil infrastructure. For example, 34% of India’s population lived in urban areas in 2021, which is increasing at 2.4% per year^8^, exacerbating pressures on already overloaded infrastructure. This is especially acute for wastewater treatment plants (WWTPs). Although functional WWTPs reduce AMR release into receiving waters^9–12^, insufficient treatment capacity results in untreated discharges that transform receiving water bodies into conduits for AMR spread, including transmission of pan-resistant strains, and emergence through mutation and HGT^13,14^. Despite the importance of quantitative source apportionment, it has remained rare in LMIC settings, limiting our ability to prioritize intervention strategies based on evidence.

The consequences of inadequate waste management and AMR spread are exemplified by conditions in the Musi River. The Musi has significant economic and societal importance^15^, but has also become a major waste disposal site for municipal and industrial wastewater from the growing city of Hyderabad, Telangana (Southern India). Importantly, Hyderabad is a global pharmaceutical manufacturing hub, infamous for high antibiotic pollution^16–19^. The construction of common effluent treatment plants (CETP) for industrial effluents and a zero liquid discharge policy for industries offers an opportunity to assess drivers of AMR spread in the Musi River, which is key to prioritising remedial actions to reduce AMR exposures across the watershed.

The Musi serves as a good model for assessing AMR fate and transport in urban river catchments in regions with warm climates (enabling rapid microbial turnover), insufficient WWTP capacity, and low dilution ratios where river discharges are small relative to waste loading^10,20–22^. These conditions are representative of many rivers including several studied rivers such as the Code, Indonesia^23^, Bagmati, Nepal^24^, Brisbane, Australia^25^, Virilla, Costa Rica^26^ and the Indian rivers Sabarmati^27^, Mutha^28^ and Kshipra^29^. Nevertheless, past studies have rarely explained spatiotemporal variation of AMR levels in terms of point-source discharges and mass loadings using hydraulic modelling approaches. Such quantitative frameworks are crucial for distinguishing levels of contribution from various industrial and municipal inputs.

Here, we integrated field monitoring and hydraulic modelling to quantify drivers of AMR in the Musi River running through Hyderabad city. Water and sediment samples were collected at 10 locations along different parts of the catchment spanning 153 km and during the wet and dry season to assess how spatial and seasonal factors impact *in situ* AMR. The objectives were to (1) characterise the spatial and seasonal distribution of AMR determinants and physicochemical conditions required for mathematical modelling and risk analysis, (2) assess correlations between the different AMR determinants and with environmental factors, (3) quantify contributions from municipal versus industrial point-sources through hydraulic modelling to explain observed patterns, and (4) identify possible water quality proxies for cost-effective AMR hotspot detection to guide interventions and prioritise sites for more detailed AMR monitoring and analyses.

## 2. Results and discussion

### 2.1. Spatio-temporal patterns of environmental conditions and AMR determinants along the river

Gradients of environmental conditions were marked. All analysed water quality (WQ) parameters followed a distinct spatial gradient along the Musi River (**Fig. S1**). Water from city sites was severely contaminated across seasons, with high concentrations (mean±SD, mg/L) of COD (168±97), TOC (14±5), NH_3_-N (62±55), TN (26±13) and TP (26±17), whereas DO remained near-anoxic (<1 mg/L). Several WQ parameters (pH, EC, COD, etc.) exceeded permissible limits set by India’s Central Pollution Control Board (CPCB) within the city, classifying the water as ‘very poor’^30^. Pollution was naturally attenuated at downstream sites (COD 74±29; DO 5±1), though levels remained altered relative to the upstream site (COD 37 ±4; DO 7±2) (**Fig. S1**). In contrast, only some of the corresponding sediment physicochemical conditions showed distinct gradients, with conductivity, ammonia and organic matter elevated in the city (**Fig. S2**).

Abundances of antibiotic resistance and other genes showed distinct patterns along the river stretches. Seven ARGs and the *intI1, uidA* and 16S rDNA genes were quantified in river water (**Fig. 1**) and sediment samples (**Fig. S3**) from the wet and dry season. The most abundant genes (log_10_ copies/mL) in city water samples for both seasons were the 16S rDNA (7.81±0.81), followed by *intI1* (6.53±0.96), *sul2* (6.17±0.82), *aph(3”)-Ib* (6.11±1.09)*, tetW* (5.56±1.06), *ermF* (5.55±1.53) and *qnrS* (5.03±1.09). Conversely, *bla*_CTX-M_ (3.97±0.84), *bla*_NDM_ (3.72±0.84), and *uidA* (3.35±1.24) concentrations were lowest in both seasons. The abundances of all ten genes followed consistent spatial gradients similar to WQ, with the lowest levels found at the upstream site, highest within the city, dropping to varying extents downstream (**Fig. 1**). During the wet season, water column gene levels increased more gradually as water passed through the city and also declined more gradually downstream. The ARG abundances in the city sites were substantially higher than those in anthropogenically-impacted European rivers, such as the Suze River in Switzerland^31^ and the Danube River in the most polluted stretches in Central Serbia and the tributary Arges river in Romania^32,33^, but comparable to concentrations in raw municipal wastewater in Hyderabad^12,34,35^.

**Fig. 1.**
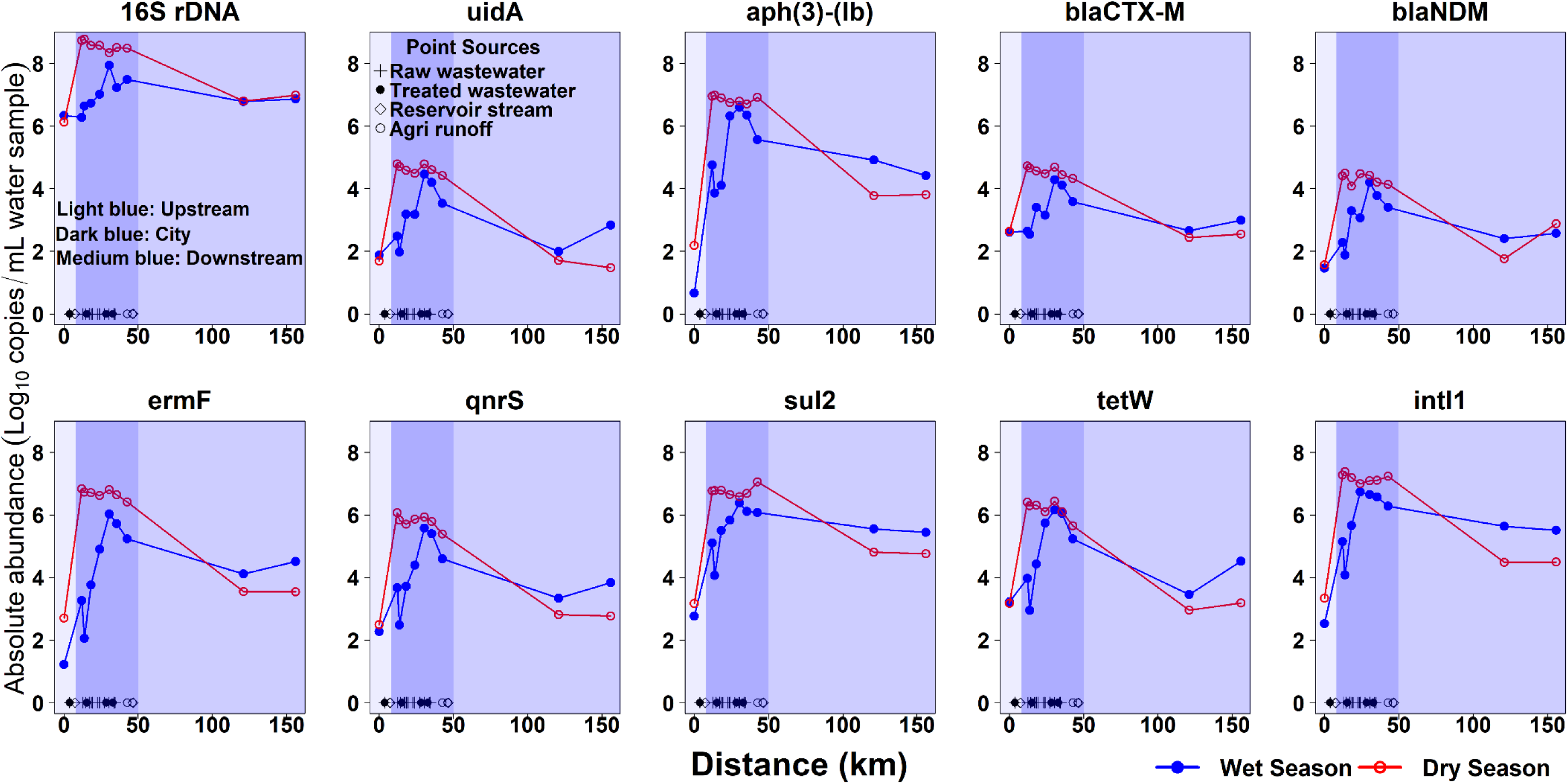
Absolute abundance of genes along the Musi River in dry (red) and wet season (blue) water samples. All panels have the same scale showing log transformed absolute abundances of 16S rDNA, *uidA*, *intI1* and seven ARGs. The river can be divided into three stretches: (a) upstream (light blue), (b) within the city (dark blue) and (c) downstream of the city (medium blue). Point-sources of raw wastewater, treated wastewater, reservoir flows, and agricultural runoff are indicated by symbols. See **Fig. S3** for equivalent sediment sample data.

Unlike water samples, no consistent seasonal patterns in gene levels were evident in the sediment samples, although concentrations were generally higher, albeit more variable, within the city (**Fig. S3**). The high concentrations in the sediments suggest that sediment acts as an important ARG reservoir, accumulating ARGs from the point-sources under low-flow conditions during the dry season. Overall, the genes analysed in water and sediment columns indicated that the Musi in Hyderabad was highly polluted with ARGs in both seasons, although spatial patterns of detected genes differed between sediment and water column samples. On a positive note, ARGs declined downstream of the city, suggesting natural recovery was rapid once pollution dropped.

Bacterial abundances showed distinct spatial and seasonal patterns that were consistent with gene patterns. ‘Total’ Gram-negative heterotrophic bacteria (OSS), *E. coli* (TBXB), and ESBL- and CARB-resistant bacteria were quantified in water and sediment samples. In general, incubation at 30℃ or 44℃ did not alter spatio-temporal distribution patterns of total GN-heterotrophic bacteria or *E. coli* in the water samples (**Fig. S4**; see **SI Section 1.1** for details). That is, viable bacterial concentrations (log_10_ CFU/mL, mean of OSS30, OSS44 and TBXB44) were lower upstream (0.46), much higher in the city (4.88±0.49), and lower again downstream (2.15±1.17) in the dry season. These spatial patterns were similar to those observed for ARGs indicating consistent pollution sources driving both bacterial and resistance gene abundances. In contrast to the dry season, bacterial abundances were elevated everywhere in the wet season, similar to levels in dry season city samples (**Fig. S4A**). This pattern aligns with studies of the Ganga river, India^36^ and parts of the Danube river, Europe^33^, where municipal wastewater discharges drive distinct spatial patterns of bacterial contamination.

The pattern for ESBL-resistant bacteria was also similar, with abundance below detection limit (BDL) upstream, higher in the city (log_10_ 4.54±0.63 CFU/mL, mean of ESBL30 and ESBL44), and returning to BDL downstream in the dry season (**Fig. S5A**). In the wet season, levels in the city were similar to those found within the city during the dry season. Conversely, CARB-resistant bacteria were BDL in the city (and elsewhere) in the dry season, while the wet season pattern reflected ESBL. The consistently higher ESBL and CARB counts in the wet season (**Fig. S5A**) suggest elevated flows reduce sedimentation of water column bacteria and/or increase scouring of microorganisms previously deposited in sediments, which can occur in rivers in monsoonal climates^37^.

Sediment bacterial abundances varied seasonally, similar to water samples and were generally higher in the wet season than in the dry season across all media (**Fig. S4B**) (for values see **Supplementary Data 1**). In contrast to the distinct spatial gradients observed in water samples, sediments showed no discernible upstream to downstream patterns during the dry season. During the wet season, abundances were slightly higher in the city and downstream sediments compared with upstream, though these differences were less dramatic than those in the water samples. ESBL- and CARB-resistant GN heterotrophs were with one exception all BDL in sediments in the dry season, while they were all at similarly higher levels in the wet season (**Fig. S5B**). This seasonal dichotomy of bacterial abundance suggests that increased wet season flows cause sediment resuspension, mobilising sediment-associated resistant bacteria into the water column, while concurrent urban runoff contributes additional microbial inputs to the river^38^.

Five hypotheses were tested regarding relationships between viable counts (per volume) at different conditions and viable counts versus gene counts quantified by qPCR: (i) counts at 44℃ would correlate with, but be less than, counts at 30℃ on the same medium; (ii) counts on CARB media would correlate with, but be less than, counts on ESBL media; (iii) counts on ESBL media would correlate with *E. coli* counts because both relate to levels of municipal wastewater pollution; (iv) counts on OSS media would best correlate with 16S rDNA gene counts; and (v) counts of *E. coli* would correlate with *uidA* gene counts. Data were consistent with all five hypotheses, but no firm conclusions could be made because of the highly scattered viable count data (**Figs. S6, S7, S8** and **Table S1**). The gene data were clearly more consistent across samples and therefore used in all subsequent numerical analyses.

### 2.2. Multivariate analysis of patterns of AMR determinants and environmental conditions

Visualizing gene abundances, bacterial counts, and water quality parameters indicated that the Musi catchment can be divided into three stretches: upstream, city, and downstream (**Figs. 1, S1, S4**). To objectively validate these patterns and test for statistical significance, hierarchical cluster analysis (HCA), principal component analysis (PCA), and permutational multivariate analysis of variance (PERMANOVA) were employed.

Hierarchical clustering of water samples revealed season-dependent spatial patterns. In the dry season, city sites formed a distinct cluster, whereas upstream and downstream sites clustered together (**Fig. 2A**). Conversely, in the wet season, city sites split with late-city sites remaining a distinct cluster, but early-city sites now clustering with downstream samples—the upstream site remaining distinct (**Fig. 2B**). This in-city split suggests the city stretch contains two sub-stretches in the wet season that merge into a single pollution zone during the dry season. These clustering patterns were based on UPGMA clustering with Euclidean distance, but similar clusters were obtained for all four combinations (UPGMA or Ward’s clustering combined with Euclidean distance or Bray-Curtis dissimilarity), confirming the robustness of the results (see **Supplementary Data 2**). PCA corroborated these patterns, showing a tight cluster of all city sites in the dry season and a more spread and mixed pattern in the wet season (**Fig. 2C-F**). All seven ARG vectors pointed in similar directions in PCA ordination space, indicating similar AMR pollution sources in the river.

**Fig. 2.**
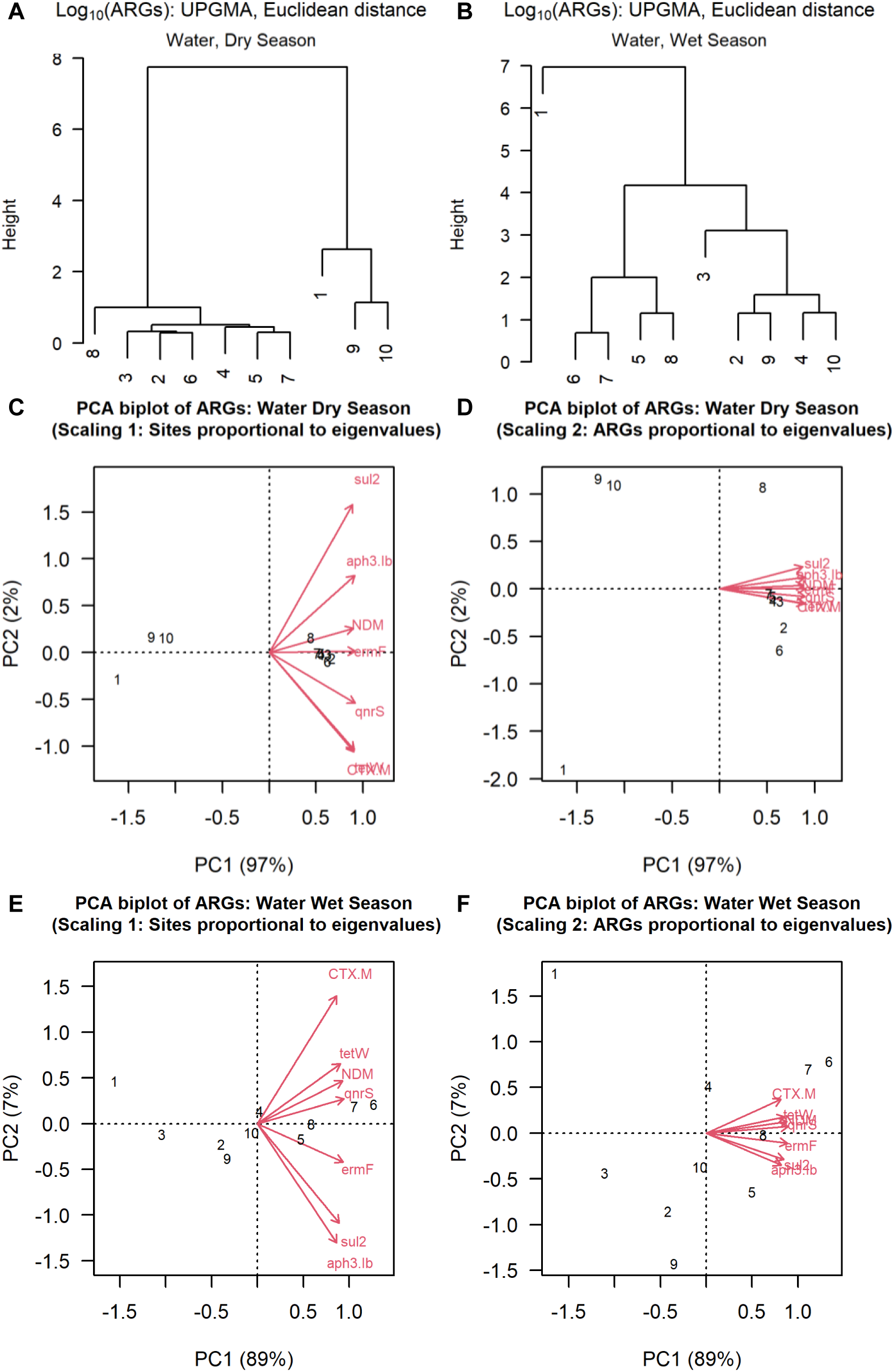
Hierarchical clustering and ordination of water samples comparing dry and wet seasons based on Log_10_ transformed absolute abundances of seven ARGs. Sampling locations are numbered along the river: upstream (1), city (2-8) and downstream (9-10). **(A-B)** Hierarchical clustering (UPGMA) of Euclidean distances shows the separation of the city from up- and downstream samples in the dry season, whereas only late city sites cluster in the wet season. **(C-F)** PCA biplots for dry **(C-D)** versus wet **(E-F)** seasons. Scaling 1 correctly represents sites with their ARG profiles while Scaling 2 correctly represents genes (vectors).

Since the sampling design was not balanced (one upstream, seven city, and two downstream sites), and variables were not independent and normally distributed, PERMANOVA (adonis2 from the vegan package) was used to test for significant spatial and seasonal differences. Both stretch and season were significantly different (*p* < 0.05) for most water column variables (**Table S2**). Stretch was significant for all 10 genes, six of seven WQ parameters (except NO_2_^−^), and all bacterial counts, *i.e*., 19 of 20 parameters (ARB data were insufficient for analysis). Season was significant for nine of 10 genes (except *tetW*), five of seven WQ parameters (except DO and NO^−^), and all bacterial counts, *i.e*., 17 of 20 variables (see **SI Section 1.2** for details). These results confirm that both spatial location and seasonal flow characteristics significantly structure AMR determinant distributions in the Musi water.

In contrast to water sites, sediment sites showed weaker spatial and temporal differentiation. Stretch was significant for only five of 10 genes (*intI1, aph(3”)-Ib, ermF, qnrS, tetW*), the sum of ARGs, two of seven SQ parameters (total nitrogen and conductivity), and two of three bacterial counts, *i.e*., nine of 20 parameters in total. Season was significant for only two genes (*uidA*, *bla*_CTX-M_), but six of seven SQ parameters, and all bacterial counts (**Table S2;** see **SI Section 1.2** for details). The attenuated spatial differentiation in sediments likely reflects long-term accumulation coupled with transport of ARGs and ARBs, which obscures the longitudinal gradients observed in water samples. Thus, sediments function as integrative archives of historical contamination rather than reflecting instantaneous pollution dynamics^38^.

Water ARG concentrations were scaled to sediment concentrations to enable comparisons between the two environmental compartments. Gene concentrations were expressed as gene copies per mL for water and gene copies per mg of organic matter (OM) for sediment samples to account for differences in sample matrices (see **Supplementary Data 2** for scaling details). Sediment gene concentrations were expressed relative to organic matter rather than dry mass because inorganic particles such as sand contributed to dry mass but were inaccessible to bacteria and their genes, and inorganic content varied substantially across sites.

These scaled data revealed season-dependent coupling between water and sediment ARG profiles (**Fig. 3**). In the wet season, sediment and water samples clustered separately, suggesting higher river flows decoupled their ARG content. This decoupling likely reflects increased lateral dispersion in the water column and increased sediment resuspension combined with reduced exchange between consolidated sediments and overlying water. Conversely, dry season clustering showed selective coupling: city water samples remained distinct, while upstream and downstream water samples clustered to some extent with their corresponding sediments. Inner-city sediment samples formed separate sub-clusters in both seasons, indicating persistent, high-level AMR contamination year-round in central Hyderabad. This seasonal shift suggests spatially homogeneous AMR pollution during low-flow conditions, with spatial differentiation emerging during elevated flows as it takes more point-sources to pollute the river after entering the city and then spreading further downstream. Further interpretations of these multivariate patterns and their connections to linear discriminant analysis are provided in **Section 2.4 (SI Section 1.2)**.

**Fig. 3.**
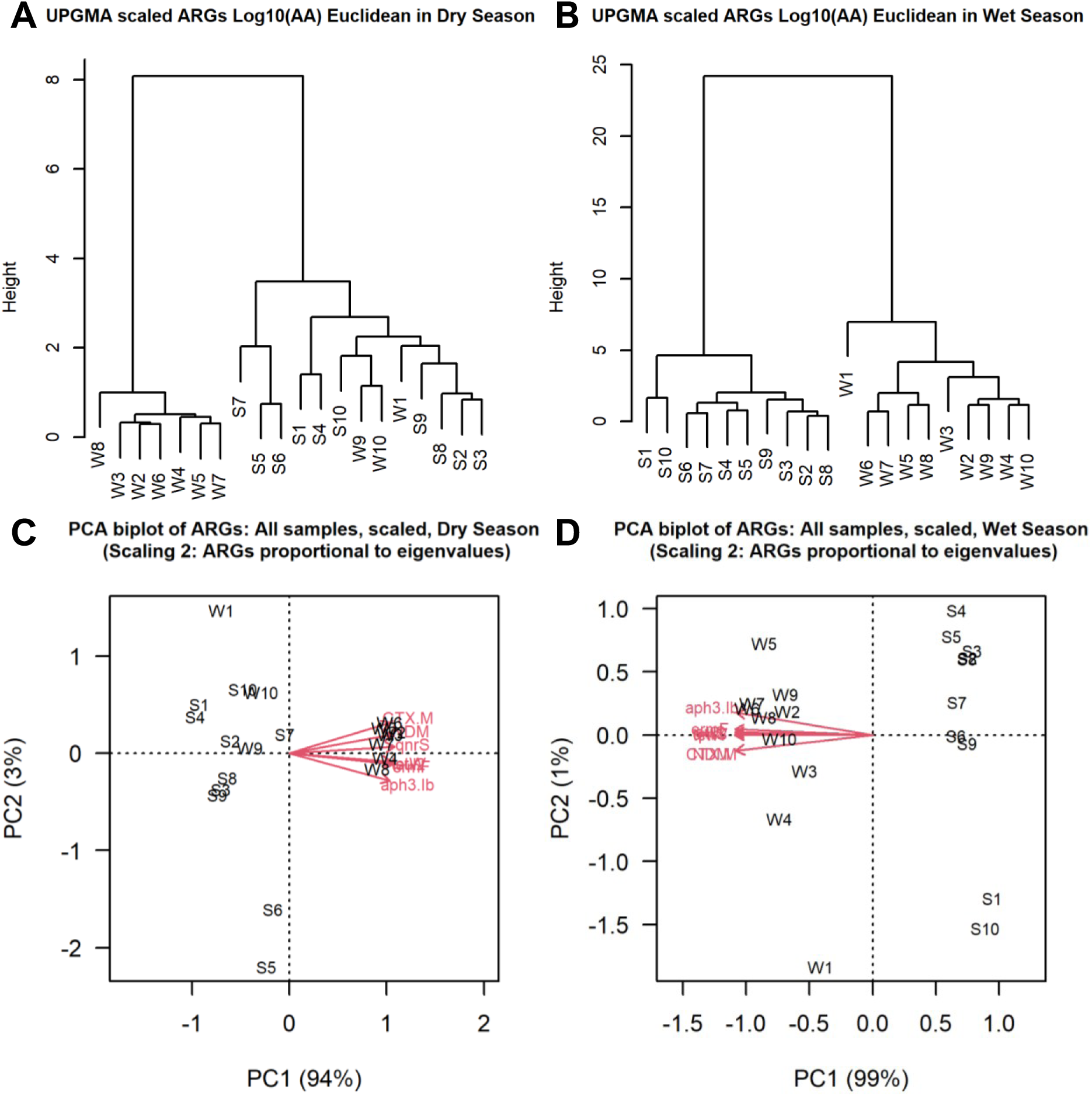
Joint clustering of sediment and water samples along the Musi in the wet and dry seasons. Sampling locations are numbered along the river: upstream (1), city (2-8) and downstream (9-10), while W represents water and S represents sediment samples. This joint analysis of water and sediment samples is based on Log_10_-transformed absolute abundances of ARGs. Since water and sediment ARG concentrations had a different basis, data were scaled to make them more comparable. (**A-B**) Joint hierarchical clustering of the Euclidean distances between ARGs, either in the dry (**A**) or wet season (**B**). (**C-D**) PCA biplots, either in the dry (**C**) or wet (**D**) season. These biplots used Scaling 2, which correctly depicts genes (vectors).

### 2.3. Source apportionment of AMR determinants in the Musi

Hydraulic modelling explained the observed spatial patterns of ARGs and ARBs in the Musi. We identified and quantified 25 point-sources of wastewater and other inputs within the area of investigation and calculated mass loadings using discharge estimates and a hypothetical conservative tracer (**Supplementary Data 3**). Point-source discharges were quantified using operational records for WWTP discharges and population-based estimates for raw wastewater production using population density and per capita wastewater production rates (150±25 L/d) from local government and other sources (see **SI Section 1.3**). The hypothetical inert tracer concentration was set to 1.0 for untreated sewage and 0.16 for treated WWTP effluent, parameterized based on measured COD remaining in WWTP effluents and consistent with what remains of volatile solids (12%), 16S rDNA (19%), and total ARGs (22%) in effluents from the four main WWTPs along the Musi^12^. Details of point-sources, sampling sites and discharge, tracer, and loading calculations are provided in **Supplementary Data 3**.

Mass balance calculations for the tracer suggested that 60-80% of river water in the city stretch was derived from raw wastewater inputs in the dry season, with total bacterial abundance (16S rDNA) ranging from 2-6×10^8^ copies/mL (**Figs. 4A, S9**). Sharp increases in bacterial abundance coincided spatially with tracer spikes at point-source discharge locations, validating the mass balance approach. In contrast, during the wet season, monsoon runoff and increased discharges released from Osman Sagar and Himayat Sagar reservoirs provided higher dilution, reducing raw wastewater contributions to 20-40% of river flow and decreasing bacterial abundance by an average of 1.52 log_10_ (0.19-8×10^7^ copies/mL) compared to the dry season (**Figs. 4B, S9**). Importantly, WWTP discharge points (tracer = 0.16) led to localized decreases in bacterial abundance and ARG levels, demonstrating that WWTPs improved local water quality (**Fig. 4**). This is in stark contrast to countries where the bulk of the wastewater is treated before entering rivers, so WWTPs pollute the rivers rather than making them cleaner as in the case of the Musi^31,32,39^.

**Fig. 4.**
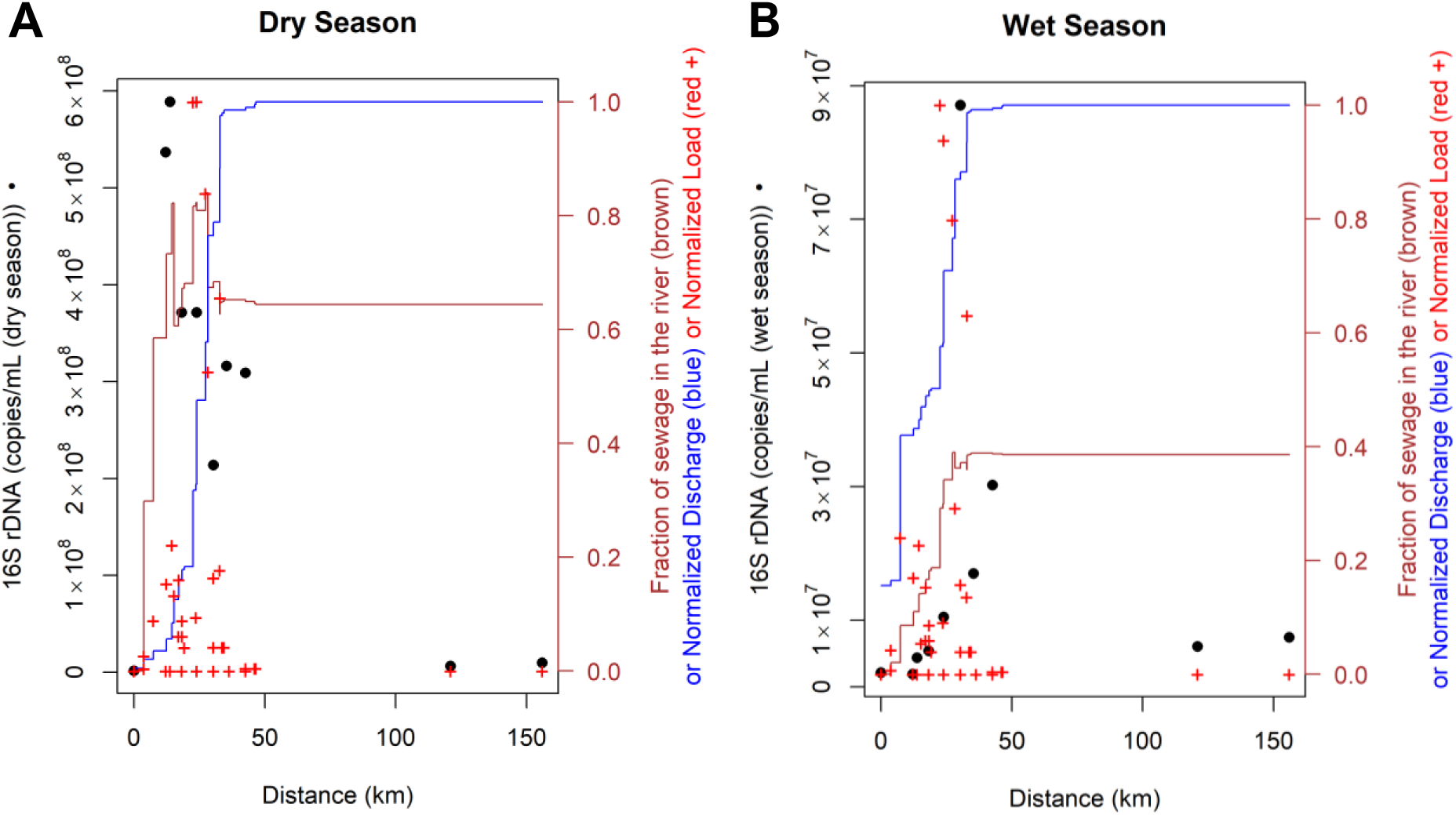
Direct comparison of the absolute abundance of the 16S gene in the river with the estimated wastewater derived fraction in both dry and wet seasons. The hypothetical tracer concentration was set to 1 in raw wastewater and to 0.16 in treated wastewater to track wastewater loads. The tracer was assumed to be inert in contrast to 16S rDNA levels (•), which declined along the river. Normalized discharge (blue stairs), normalized loadings (+) and the fraction of wastewater in the river based on tracer mass balance calculations (brown stairs).

Predicted spikes of the tracer and untransformed 16S rDNA gene abundances (**Fig. 4A**) both indicated that bacterial levels (which mirror ARGs and other measured variables) increased sharply at point-source locations. However, 16S abundances declined progressively downstream of these inputs, in contrast to the tracer that was assumed to be inert. The downstream decline of the 16S abundance was likely due to microbial die-off, which was not quantified and may vary with environmental conditions. Nevertheless, the tracer calculations allow the quantification of municipal wastewater inputs into the river. Interestingly, the observed data imply that the Musi River itself acts as a “WWTP” for the city’s wastewater, treating approximately 59% of Hyderabad’s sewage during both the dry and wet seasons (see **Supplementary Data 3**). A similar phenomenon was seen in community sewer lines in an European study on the fate of AMR in several urban catchments^40^. However, the ecological consequences and health risks of this wastewater dominance are severe as city stretch waters caused 100% mortality in zebrafish (*Danio rerio*) embryos within 96 hours post fertilization, compared to 0% mortality when exposed to upstream water^41^, indicating that the Musi River has been transformed from a viable aquatic ecosystem into a *de facto* wastewater conveyance channel.

These mass balance calculations show the importance of considering point-source mass loadings (*i.e*., mass transferred per time), rather than just concentrations, when developing practical solutions to problems like AMR spread in rivers. The current “[WHO Guidance on waste and wastewater management in pharmaceutical manufacturing with emphasis on antibiotic production](https://www.who.int/news/item/21-12-2023-public-consultation-on-antimicrobial-manufacturing-effluent-guidance)” is based on point-source concentrations, assuming a fixed dilution factor of 10 upon entry into a river. This concentration-focused approach creates a regulatory loophole as facilities can achieve compliance through pre-discharge dilution without reducing actual pollutant mass inputs. Such a loophole can be addressed by also stipulating load limits in environmental standards, especially when considering AMR genes and bacteria released into the environment.

Pharmaceutical manufacturing contributed far less to the AMR burden in the Musi than municipal wastewater. Given that India is the world’s third-largest pharmaceutical producer with over 400 rivers and approximately 2.4 million other freshwater bodies^42^, the country faces dual AMR pollution risks. Two questions guided this analysis: (1) How typical is the Musi of Indian rivers? and (2) Do manufacturing waste inputs alter AMR in the river?

Regarding the first question, rapid urbanisation in India has greatly outpaced the construction of new WWTPs and supporting sewerage infrastructure. The “National Inventory of Sewage Treatment Plants” of India^43^ indicates that only 28% of municipal wastewater is treated nationwide and only 17% in WWTPs compliant with discharge standards. Hyderabad treats ∼41% of its wastewater (see **Supplementary Data 3**), above national average, yet ∼59% of municipal wastewater remains untreated and is discharged directly into the environment. This infrastructure deficit makes the Musi representative of many Indian rivers where treatment capacity lags behind population growth, a similar pattern also observed globally in rapidly urbanising countries^44^.

Regarding the second question, a distinctive aspect of the Hyderabad sewershed is that it has extensive antimicrobial manufacturing and processing units, which discharged wastewater into the local sewer drains until the construction of two treatment plants (see below). Thus, Hyderabad has been infamous for pollution from manufacturing since Larsson et al.^16^ in 2007 found ciprofloxacin concentrations in surface waters near the city of up to 31 mg/L. Subsequently, ciprofloxacin concentrations as high as 5.5 mg/L were found in the Musi during the dry season^19^. These concentrations were measured after the construction of two Common Effluent Treatment Plants (JETL in 1989 and PETL in 1995), designed specifically for the antimicrobial manufacturing cluster, which discharge their treated effluents into the Amberpet municipal WWTP for additional treatment and dilution before reaching the Musi River, indicating ongoing illegal dumping. Recent studies found much lower concentrations of antimicrobials in surface waters in the region, suggesting stricter enforcement has reduced illegal discharges into the Musi^17,18^. For example, ciprofloxacin levels were much lower than previous observations, *i.e*., only 19.3 μg/L in local sewer drains and 7.28 μg/L in local industrial effluents and much lower in the Musi water (**Table S3**). Based on our mass-balance calculations, the PETL/JETL discharge becomes diluted upon entering Amberpet WWTP and again upon entering the Musi to the extent that only 4% of the river water after Amberpet WWTP derives from PETL/JETL discharge in the dry season (dilution factor of 27), and these calculations did not consider treatment and other losses along the sewer flow path (see **Supplementary Data 3**). Therefore, the manufacturing waste inputs and impacts on the Musi River appear to be small relative to municipal inputs at the catchment scale. Previous studies detected several antibiotics and ARGs (*aph(3”)-Ib*, *ermF*, *qnrS*, *sul2*, *bla*_CTX-M_, *bla*_NDM_) in drains of Hyderabad^34,35^, the same genes were abundant in the Musi as expected from local antibiotic sales patterns in Hyderabad (**Fig. S10**). However, the spatial concordance between ARG levels and municipal drain inputs (**Figs. 1, 6**), combined with volumetric dominance, demonstrates that nowadays untreated municipal wastewater drives AMR pollution in the Musi River at the catchment scale.

### 2.4. Water quality parameters as proxies to predict AMR hotspots

Having established that municipal wastewater drives AMR pollution in the Musi (**Section 2.3**), we investigated whether simple water quality parameters could serve as cost-effective proxies for identifying AMR hotspots, enabling rapid screening in resource-limited settings. Water quality indicators (DO, COD, TOC, TN, etc.) exhibited spatial patterns mirroring ARG and bacterial distributions: low pollution upstream, very high in the city, and moderate downstream (**Fig. S1;** detailed explanation in **SI Section 1.4**). This concordance implied wastewater pollution as the common cause for water quality degradation and AMR prevalence, suggesting that traditional water quality indicators may serve as surrogates for AMR contamination.

To quantitatively test whether water quality parameters could discriminate river stretches across seasons and predict AMR prevalence, Linear Discriminant Analysis (LDA) was performed on z-score standardised water quality (WQ) data (**Fig. S11**). LDA is a constrained ordination method to find linear combinations of explanatory variables that best separate predefined categories, here river stretches distinguished by their significantly different ARG profiles. Our full LDA model using all seven non-collinear WQ parameters (WQ7: pH, DO, COD, NO_2_^−^, NO_3_^−^, NH_4_^+^, TN) clearly discriminated between stretches and seasons with 100% classification accuracy (**Fig. 5A**). To develop the simplest field deployable model, parameters were removed from the full model in stages (since there were 127 possible combinations of the seven parameters). By removing variables individually, two were found to have little effect, resulting in a model with four variables (WQ4: pH, DO, COD, TN) with 95% accuracy that maintained clear stretch separation and was logical in terms of river water chemistry and microbiology (**Fig. 5B**). Further simplification demonstrated that the two-parameter model combining DO and TN achieved 90% discrimination accuracy, superior to all other two-parameter combinations, representing a good balance between analytical simplicity and predictive power (**Fig. 5C**). Note that despite inaccuracies in classifying stretch and season, the distinction polluted versus less-polluted was fully accurate. Although the DO+TN model was less accurate than WQ4, it could provide a practical, cost-effective tool for rapid AMR hotspot identification in resource-limited settings.

**Fig. 5.**
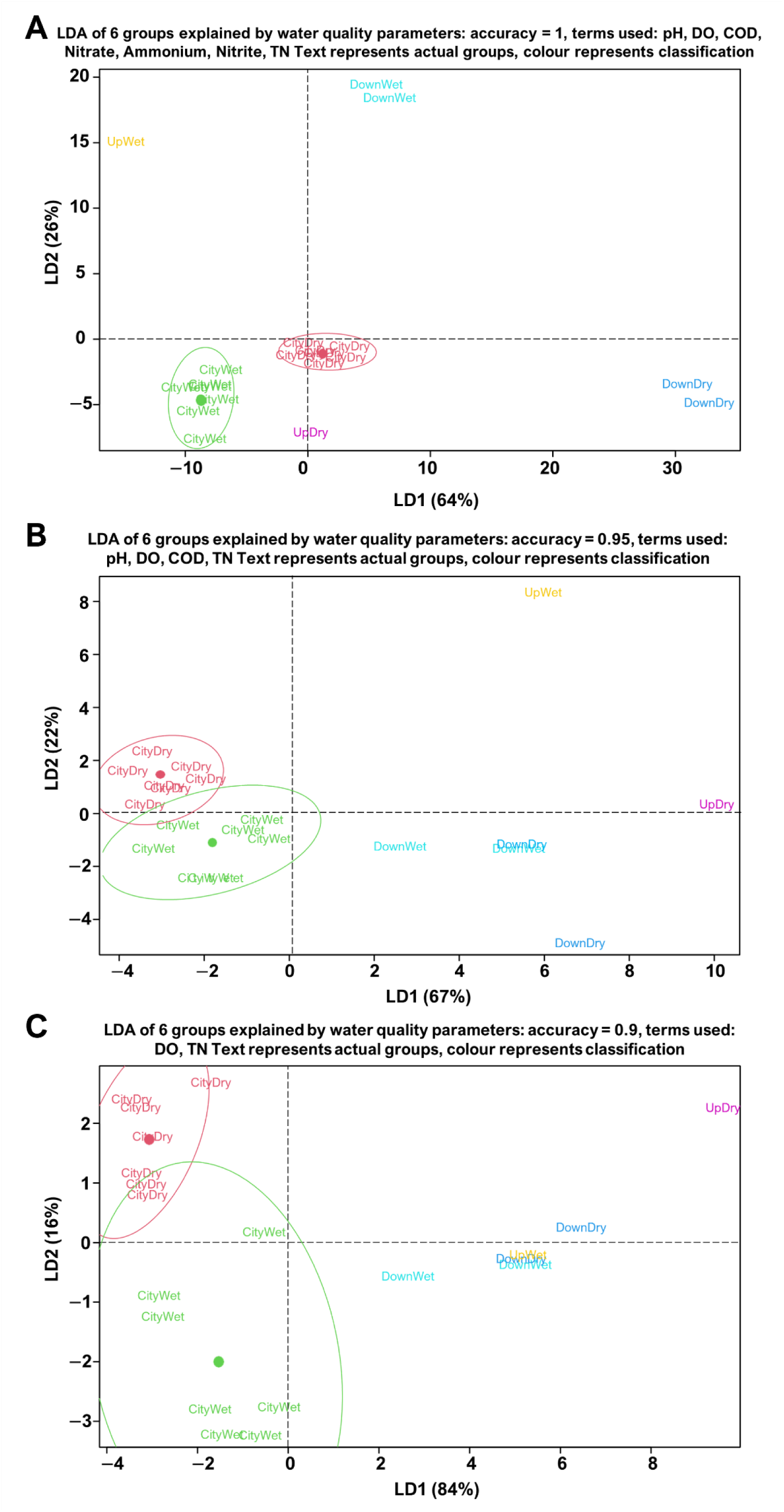
Linear discriminant analysis (LDA) showing that the three river stretches identified by clustering and ordination of ARGs can be discriminated by environmental characteristics in both seasons (resulting in 6 groups). Text labels combine actual stretch (Up, City, Down) and season (Dry, Wet) while colours indicate the LDA classifications of the sites, which are mostly consistent (accuracy of classification 90%-100%). Ellipses show 95% confidence intervals. **(A)** Discrimination using the full set of seven independent water quality parameters (WQ7). **(B)** Discrimination without the N-cycling ammonium, nitrite and nitrate (WQ4). **(C)** Discrimination with only DO and TN as water quality parameters (DO plus TN).

Ott et al.^45^ previously proposed the use of DO and ammonia as proxies for AMR pollution based on the Skudai River, Malaysia. Applying their DO+ammonia model to the Musi data provided similar results to our DO+TN model, while the latter proved better for the Musi data (**Fig. S12**). This difference reflects distinct nitrogen cycling dynamics between these rivers. which can explain why TN provided a more reliable proxy for AMR hotspots, particularly in rivers with strong redox gradients such as the Musi. The superiority of TN over individual nitrogen species (ammonia, nitrite, nitrate) warrants a mechanistic explanation especially as this has practical implications for monitoring diverse aquatic environments. Wastewater inputs introduce organic carbon and nitrogen while stimulating microbial respiration, which depletes DO. In the Musi, DO declined to near-anoxic levels (<1 mg/L) in the city stretch during the dry season, creating conditions for denitrification that consumed residual nitrite and nitrate (**Fig. S1**). Downstream, reduced wastewater loading and re-oxygenation increased DO, halting denitrification and enabling nitrification, which oxidised ammonia to nitrite and nitrate. This caused transient spikes in oxidized nitrogen species at the downstream site 9 (**Fig. S1**) despite declining wastewater influence. Consequently, ammonia, nitrite, and nitrate concentrations fluctuated along the river dependent on the presence of oxygen, making each individual species an unreliable indicator of total wastewater loading. In contrast, TN integrates most nitrogen forms and is conserved during several of these transformations, providing a more stable proxy for cumulative wastewater inputs that is less dependent on local redox conditions. PERMANOVA confirmed TN significantly distinguished river stretches (*p* < 0.001) (**Table S2**). This explains why Ott et al.^45^ found ammonia to be a reliable proxy as the Skudai River remains oxygenated with low N-cycling, so ammonia concentrations directly reflect wastewater inputs. However, when DO varies like in the Musi, representative of many urban rivers in LMICs receiving high organic loads, nitrogen speciation varies dynamically, suggesting DO+TN could be a more robust proxy for wastewater pollution and associated AMR.

Limitations. Although EMBRACE-WATERS guidelines for studying AMR in wastewater and aquatic environments were followed^46^, our study had sampling limitations and analytical challenges. Downstream sampling sites were too far away to be sampled on the same day. Resource limitations restricted sampling to once per season, precluding a more extensive temporal analysis. The study faced three analytical challenges. First, variability in sediment compositions, ranging from mineral-rich (sandy) to organic-rich samples, complicated comparisons between samples. We addressed this by normalising concentrations to the organic fraction (organic matter) of the sediments. This approach, while not ideal, was deemed the best available option. Further research across diverse sediment types is necessary to refine this method for future updates of the EMBRACE-WATERS guidelines. Second, because of using organic matter as denominator for concentrations in sediment samples, comparisons to water samples with volume as denominator were not straightforward. To facilitate this comparison, ARG concentrations in water samples were scaled to align with average sediment concentrations, but this is unlikely to have made water and sediment concentrations fully comparable. Third, correlations between viable bacterial counts and gene copy numbers from qPCR were poor due to large scatter in count data, likely due to differences in the sample preparation methods: larger water volumes (∼200 mL) for qPCR should be less affected by small-scale environmental heterogeneity than the diluted 100 μL samples for viable counts. Moreover, viable counts will be affected by variable detachment of cells from particle surfaces during sample homogenisation while filtration for DNA extraction would not be affected. This highlights the need for methodological improvements and standardisation to enable reliable data integration across methods.

Conclusions. This study provides the first comprehensive apportionment of AMR sources in an Indian urban river by integrating field monitoring and hydraulic modelling to quantify pollution sources and thus identify practical AMR surveillance tools for rapidly urbanising regions. We studied a small river. They are not only more common than large rivers but facilitate investigation in multiple ways. Point-sources become less diluted (higher signal to noise ratio), mixing lengths are shorter and relatively clean upstream stretches, if present, are within easier reach. Mass balances suggested that, in the city stretch, 60-80% and 20-40% of river water derived from untreated municipal wastewater during dry and wet seasons, respectively. Strong correlations between predicted wastewater fractions and measured AMR determinants validated this attribution. Despite Hyderabad’s concentration of pharmaceutical industries, pharmaceutical manufacturing and processing waste discharges into the Musi likely had minimal impact on *in situ* AMR as only ∼4% of the river water following the Amberpet WWTP effluent point-source derived from industrial treatment plant effluent. Given that Hyderabad has greater than average WWTP coverage than other cities in India, it is probable that sources and conditions observed in the Musi are common in India and other countries with insufficient sewerage and wastewater treatment infrastructure. LDA showed that simple water quality markers such as DO and TN can serve as proxies for local stakeholders to identify AMR hotspots in freshwaters and to target interventions. DO and TN mirror sewage inputs, which often dominate AMR determinant loadings, particularly in rapidly urbanising countries where population growth has outpaced treatment infrastructure. These proxies are especially valuable because DO and TN are inexpensive to measure and DO is already included in current river health monitoring programmes. However, these proxies need validation in other rivers, especially when coupled with field-based methods such as DO test strips, to empower citizens to monitor and identify local AMR hotspots in their own catchments. Finally, this work highlights the value of simple hydrological and hydraulic modelling in predicting pollution loads, dilution factors, and concentrations in receiving rivers based on publicly available data. This approach identifies areas of greatest AMR exposure risk and transmission potential using mass loading data rather than concentration data from field assessments. Our methodology provides a framework for identifying and quantifying exposure pathways in future risk assessments, which are critical to better understand, prevent, and prioritise mitigation actions against AMR in places with insufficient WWTP capacities like the Musi River.

## 3. Methods

This study adhered to EMBRACE-WATERS reporting guidelines^46^ wherever possible, any deviations are indicated.

### 3.1. Study area and sample collection

The Musi is ∼260 km long and a major tributary of the Krishna River. Within Hyderabad city, it is 30-120 m wide and 2-4 m deep, depending on location and season. The average annual rainfall in its catchment is ∼800 mm, most of it falling in the monsoon or wet season (June to September). The Musi riverbed is generally rocky, consisting primarily of pink granite^47^. Two reservoirs, Osman Sagar and Himayat Sagar, exist where the Musi and Musa Rivers (a tributary of the Musi 60 km upstream of Hyderabad city) are blocked by dams, respectively. These two dams are closed most of the year except for spillway overflows during the wetter months. Therefore, flows in the Musi are often very low upstream of the city (dependent on leaky dams), and most of the flow within the city is from wastewater discharges, about 1.2 million m^3^/d of partially treated municipal wastewater, mixed with industrial wastes from across the city.

The wastewater discharged into the Musi is either untreated or released from municipal WWTPs (**Supplementary Data 3**). Two Common Effluent Treatment Plants (CETPs) for pharmaceutical and industrial waste also discharge into the Musi catchment: the Patancheru Enviro Tech Ltd. (PETL) and Jeedimetla Effluent Treatment Plant (JETL) release 1,600–2,000 and 50,000 m^3^/day, respectively. Effluents from the PETL and JETL flow through a 26 km pipeline into the Amberpet WWTP (339,000 m^3^/day), where the industrial discharges mix with untreated wastewater, before treatment and discharge into the Musi itself **(Fig. 6)**^30,48,49^. In terms of land use, the Musi catchment is both rural and urban, including many residential communities, hospitals and factories. Downstream of the city, land use includes agriculture and aquaculture, with less human impact than within the city. Sampling sites were chosen to contrast upstream water conditions with sites in the city and downstream. Since the Musi flows from a reservoir upstream of the city, only one upstream sampling point was possible. Within the city, many locations were sampled, reflecting different inputs along the river. Note that the river where we sampled may not have been fully mixed with the inputs from the more recent point-sources, despite sampling midstream from bridges, since the estimated mixing length according to the simple, generic rule of width^2^/depth^50^, ranged from 225-450 m when the river is 30 m wide to 3,600-7,200 m when the river is 130 m wide (for a depth of 2 or 4 m in both cases). Potentially incomplete mixing should have a limited effect on the results due to the large number of point-sources, with only the more recent ones potentially not fully mixed, and the larger scale separation between upstream, city and downstream stretches.

**Fig. 6.**
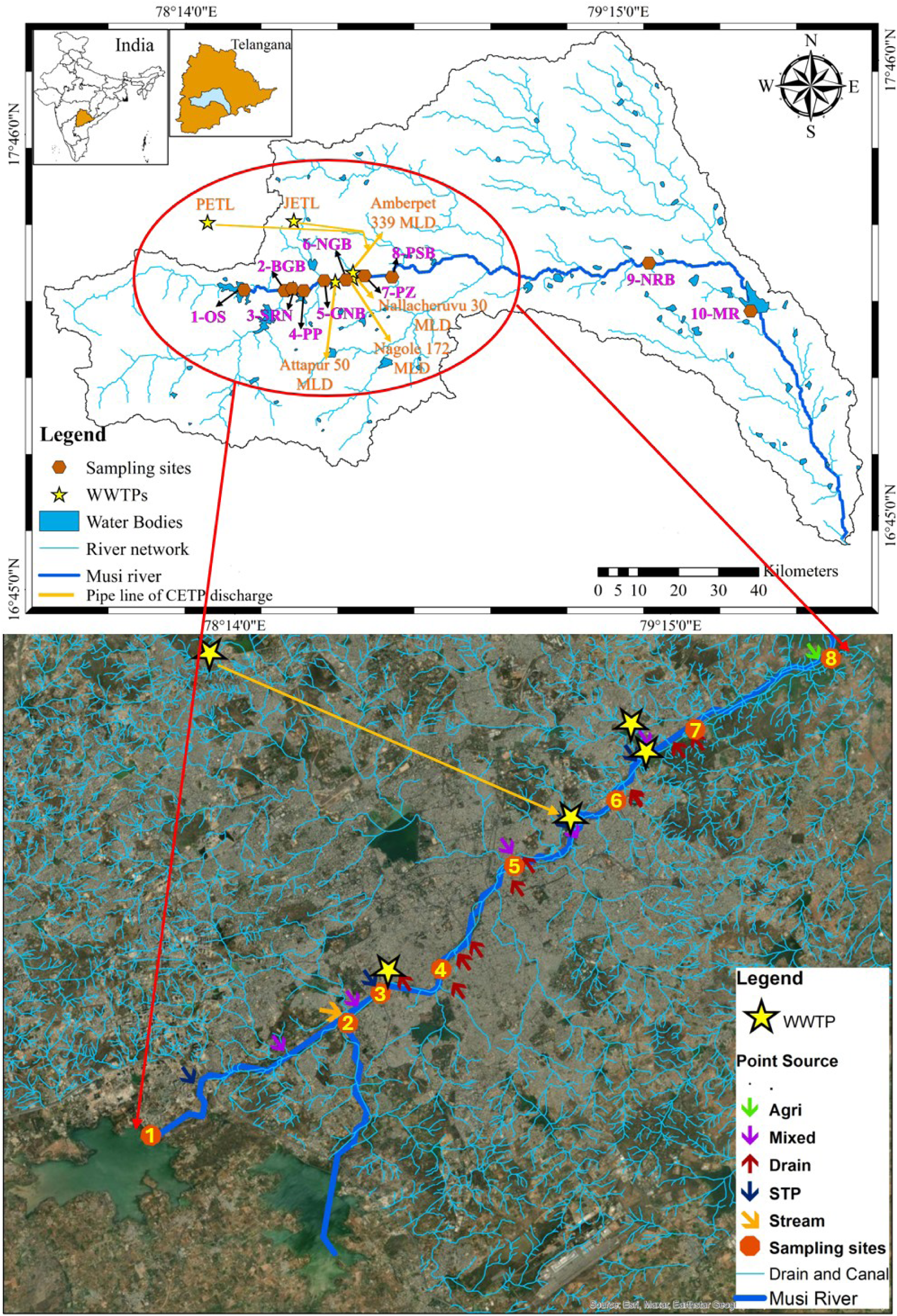
The basin of the Musi River in Telangana, Southern India, with the locations of sampling sites (numbered 1-10) and WWTP effluent discharges. A total of 25 point-sources of untreated sewage discharged through drains or treated sewage and other sources were identified within Hyderabad, between sampling points 1 and 8, see **Supplementary Data 3** for details.

Water and sediment grab samples were collected at 10 locations along the Musi River flowing through Hyderabad city, Telangana, Southern India during the wet (18-19 August 2022) and dry (11-12 March 2023) seasons **(Fig. 6**; site details in **Supplementary Data 3)**. Sampling occurred on days without rainfall for the preceding five days to minimize storm-related variability. At most sites, duplicate water samples were collected mid-stream from bridges using a bucket suspended on a rope and transferred to pre-sterilised 2-L HDPE bottles. Sub-samples were preserved immediately with H_2_SO_4_ for chemical oxygen demand (COD) analysis, whereas water temperature, pH, dissolved oxygen (DO), total dissolved solids (TDS) and conductivity were measured on-site with an Orion Star A329 portable multimeter with smart probes (ThermoFisher Scientific, USA). Duplicate sediment samples (approx. 1 kg) were collected 3 metres from the riverbank using a grab dredger (0-15 cm depth), sieved through a 2 mm sterile mesh, and transferred into pre-sterilised polypropylene bags. All samples were stored in the dark and on ice in a cooler and transported back to the lab for further processing and analysis.

### 3.2. Water and sediment quality analysis

Water and sediment physicochemical parameters were measured using standard protocols (**Table S4** and **S5**). When required for analysis (*i.e.*, organic carbon, total nitrogen (TN), ammonia, nitrate, and nitrite), water samples were filtered through 0.45 μm polyvinylidene difluoride (PVDF) membrane filters (Merck Life Sciences, Merck, India) before analysis. To establish a common basis for concentrations of bacteria and genes across sediment samples with different mineral content, dry matter (DM) and organic matter (OM) fractions were quantified using standard methods (**Table S5**).

### 3.3. Enumeration of putative *E. coli,* heterotrophic bacteria and ARBs

Seven bacterial targets were enumerated in water and sediment samples across seasons following methods described in Sonkar et al.^12^. Briefly, *E. coli* was quantified on Tryptone Bile X-β-D-glucuronide agar (TBX) (HiMedia, India) as blue-green colonies (TBXB), incubated at 44°C^51,52^, as an indicator of faecal pollution and sentinel for AMR prevalence^53^. While there is no guarantee that all blue-green colonies on TBX plates were *E. coli*, we sequenced the genomes of 58 isolates from blue-green colonies, and all were *E. coli* (Sonkar et al., in preparation), so we simply refer to putative *E. coli* as *E. coli*. ‘Total’ Gram-negative (GN) heterotrophic bacteria were enumerated on SDS-supplemented OECD synthetic sewage medium (OSS) at both 30°C for environmental and 44°C for gut-associated bacteria. Extended-spectrum β-lactamase (ESBL)-producing and carbapenem-resistant (CARB) GN heterotrophic subpopulations were enumerated on OSS supplemented with appropriate selective antibiotics^12^. These clinically relevant resistance phenotypes were selected as GN bacteria account for the majority of global AMR-attributable mortality^1^. All samples were plated in triplicate within 24 hours of collection, and colony counts recorded after 24 hours of incubation.

### 3.4. Measurement of ARGs by qPCR

DNA was extracted as previously described by Sonkar et al.^12^. The 16S rDNA gene was quantified as a proxy for total prokaryotes and to compare with culturing data. ARG targets were chosen according to Berendonk et al.^54^ and to reflect resistance to antibiotics used and produced in Hyderabad (**Fig. S12)**. Genes included fluoroquinolones (*qnrS*), aminoglycosides (*aph(3’’)-Ib*), sulfonamides (*sul2*), carbapenems (*bla*_NDM_) and third-generation cephalosporins (*bla*_CTX-M_), macrolides (*ermF*), and tetracyclines (*tetW*). Two more genes were included: *uidA* to quantify *E. coli* numbers^55,56^ and *intI1* to quantify class I integrons as a proxy for MGEs^57^. Based on preliminary analysis, DNA extracts were diluted 10-fold to reduce inhibition. The protocol was adapted from Pallares-Vega et al.^9^ as previously described in Sonkar et al.^12^. Sterile molecular-grade water was used as a no-template control (NTC) in every qPCR run for each target gene (amplification was not observed). Concentrations of forward and reverse primers, amplicon sizes of target genes and annealing temperatures are summarised in **Table S6**.

### 3.5. Quantifying pollutant point-sources along the Musi River

To quantify the drivers of AMR in the Musi River, we developed a hydraulic mass balance model to estimate river discharge and pollutant loadings from point-sources. We identified 25 point-source inputs throughout the catchment, including untreated municipal wastewater, treated WWTP effluents, agricultural runoff and mixed wastes, and estimated volumetric flow and mass loadings from each source (**Fig. S13, Supplementary Data 3**).

For each location, wastewater production rates were estimated based on local population sizes and the average municipal wastewater production rate (**Fig. S14**) specifically for Hyderabad (detailed estimates given in **SI Section 1.3, Supplementary Data 4)**. Further, flow records of WWTP and reservoir discharges were obtained. Catchment sizes and locations were based on official records and digital elevation maps. For corroboration, an alternative approach to estimate population sizes was followed, based on census data combined with satellite images to identify dwellings and allocate people to dwellings ([High Resolution Population Density Maps + Demographic Estimates](https://dataforgood.facebook.com/dfg/docs/methodology-high-resolution-population-density-maps)).

Results from the two approaches were reasonably consistent, comparing estimates using a hypothetical inert tracer that reflected wastewater inputs from point-sources. None of the measured variables could be used as an inert tracer as they were either not inert, or not exclusively tracking wastewater inputs, so our inert tracer had to be hypothetical. It was meant to reflect the typical composition of wastewater. Thus, it was parameterized based on measured values of COD in untreated (normalised to 100%) and treated wastewater (16% of untreated wastewater)^12^. Note COD was far from inert. It was chosen as a main component of wastewater that is universally measured and also because its reduction by the Musi WWTPs to 16% was similar to other relevant parameters such as VS (12%), 16S (19%) and total ARGs (22%) (**Supplementary Data 3, Sheet WWTP)**. Mass balance and loading calculations for the tracer were performed using upstream river discharge and concentration data and estimated point-source discharges to determine the relative impact of each point-source on river water quality (**Supplementary Data 3**).

### 3.6. Data processing and statistical analysis

Statistical analyses were performed with R 4.3.0 (R Core Team 2018), mainly packages ade4, vegan and MASS using the RStudio 2023.03.0 desktop (https://www.rstudio.com/) and fully documented in an R notebook (**Supplementary Data 2**). The methods and approach, including Hierarchical clustering, Principal Component Analysis (PCA), Linear Discriminant Analysis (LDA) and PERMANOVA, were guided by Borcard et al.^58^, but we are responsible for the choices made. Two clustering methods and two dissimilarity measures were compared to ensure that results were robust and not dependent on specific choices. We only report results supported by all method combinations. Ricotta and Podani^59^ was used for guidance on the choice of dissimilarity measures. Whether data should be scaled or standardised was decided after exploring and analysing the data. It was not possible to perform sample size (statistical power) calculations *a priori* due to a lack of preliminary data.

## Data availability

All data are made available as part of supplementary files.

## Code availability

All R code for all statistical analysis is made available as **Supplementary Data 2** in Rmd format.

## Supporting information

Supplementary Information (combined)

## Acknowledgements

The work of our AMRflows team has benefitted from numerous discussions with colleagues from the other four projects funded by the “India-UK Tackling AMR in the Environment from Antimicrobial Manufacturing Waste” call through the NERC-funded [Programme Coordination Team](https://indiaukamrenvironment.org/) (PCT). We would also like to thank colleagues from IIT Hyderabad for assisting with the field sampling (Jilla Adithya, Vineeth Jupaka, Sreshtha Baidya and Nikash Naorem) and Mohammed Azharuddin for his assistance with plotting the graphs in R.

## CRediT author contributions

**Vikas Sonkar:** Conceptualization, Methodology, Data curation, Investigation, Formal analysis, Validation, Visualization, Writing – original draft, review & editing. **Arun Kashyap:** Conceptualization, Methodology, Data curation, Investigation. **Rebeca Pallares-Vega:** Conceptualization, Methodology, Investigation, Formal analysis, Validation, Writing – review & editing. **Sai Sugitha Sasidharan:** Methodology, Data curation, Investigation. **Ankit Modi:** Methodology, Investigation, Data curation, Formal analysis. **Cansu Uluseker:** Methodology, Investigation, Data curation. **Sangeetha Chandrakalabai Jambu:** Writing – review & editing, Project administration. **Pranab Kumar Mohapatra:** Conceptualization, Formal analysis, Writing – review & editing, Supervision, Funding acquisition, Project administration. **Joshua Larsen:** Conceptualization, Formal analysis, Writing – review & editing, Supervision, Funding acquisition. **David W Graham:** Conceptualization, Formal analysis, Writing – original draft, review & editing, Funding acquisition. **Shashidhar Thatikonda:** Conceptualization, Methodology, Writing – review & editing, Supervision, Funding acquisition, Project administration. **Jan-Ulrich Kreft:** Conceptualization, Formal Analysis, Methodology, Writing – original draft, review & editing, Supervision, Funding acquisition, Project administration.

## Author Information

### AMRflows consortium

^1^Panagiota Adamou, ^2^Shubham Anurag, ^2^Sounak Banerjee, ^1^Ewelina Bien, ^3^Sangeetha Chandrakalabai Jambu, ^1^David W Graham, ^1^Colin Harwood, ^4^Siu Fung Stanley Ho, ^5^Rupert Hough, ^2^Chaitanya Janivara Chandregowda, ^1^Kelly Jobling, ^3^Arun Kashyap, ^4,6^Jan-Ulrich Kreft, ^2^Soumendra Nath Kuiry, ^7^Joshua Larsen, ^2^Thara Methale Velakkath Paramba, ^8^Ankit Modi, ^8^Pranab Kumar Mohapatra, ^2^Anantha Barathi Muthukrishnan, ^2^Indumathi Manivannan Nambi, ^1^Julián Osvaldo Ovis Sánchez, ^1^Rebeca Pallarés Vega, ^2^Ravikrishna Raghunathan,^2,4,6^Sasikaladevi Rathinavelu, ^3^Aravind Kumar Rengan, ^8^Wahidullah Hakim Safi, ^2^Shubhangi Sanjeev, ^2,3^Sai Sugitha Sasidharan, ^2^Vandit Kumar Shah, ^3^Vikas Sonkar, ^3^Rupali Srivastava, ^3^Shashidhar Thatikonda, ^5^Mads Troldborg, ^4,6^Cansu Uluşeker, ^9^Rama Vaidyanathan, ^4^Willem van Schaik, ^4,6^Anjali Vasudevan, ^2^Kiruthika Eswari Velmaiel, ^2^Arathy Viswanathan.

^1^School of Engineering, Newcastle University, Newcastle upon Tyne, NE1 7RU, UK

^2^Environmental & Water Resources Engineering Division, Department of Civil Engineering, Indian Institute of Technology Madras, Chennai, 600036, Tamil Nadu, India

^3^Environmental Microbiology Laboratory, Department of Civil Engineering (Environmental Engineering), Indian Institute of Technology Hyderabad, Kandi, Sangareddy, 502284, Telangana, India

^4^Institute of Microbiology and Infection, University of Birmingham, Edgbaston, Birmingham, B15 2TT, UK

^5^The James Hutton Institute, Craigiebuckler, AB15 8QH, Aberdeen, UK

^6^School of Biosciences, University of Birmingham, Edgbaston, Birmingham, B15 2TT, UK

^7^School of Geography, Earth and Environmental Sciences, University of Birmingham, Edgbaston, Birmingham, B15 2TT, UK

^8^Department of Civil Engineering, Indian Institute of Technology Gandhinagar, Gandhinagar, 382355, Gujarat, India

^9^Dr. M.G.R. Educational and Research Institute, E.V.R. Periyar Salai, Adayalampattu, Chennai, 600 095, Tamil Nadu, India

