## Supplementary Information (combined) for "Hydraulic modelling reveals untreated sewage, not pharmaceutical waste, drives antimicrobial resistance in a small river running through a big city"

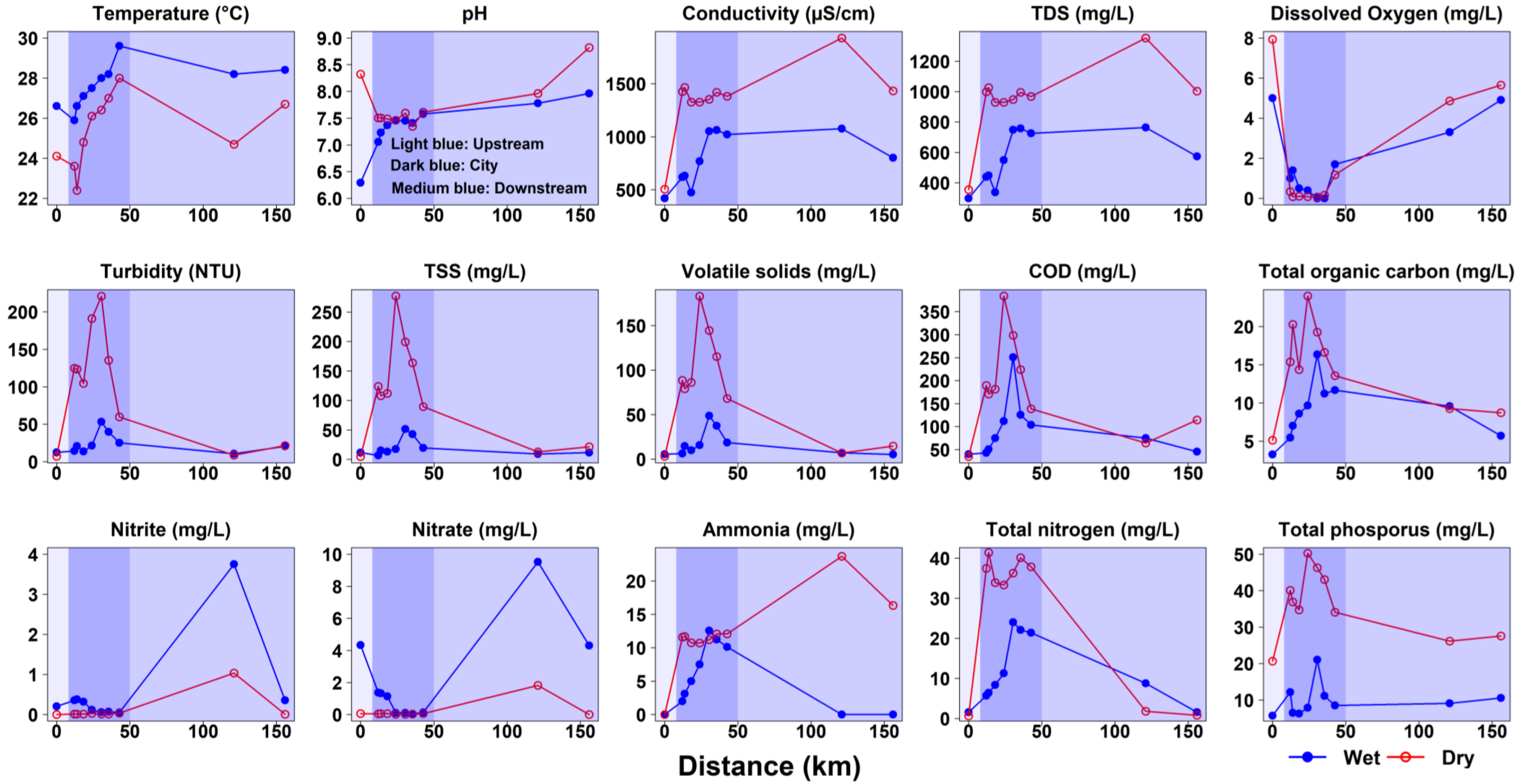


**Fig. S1. Water samples: All analysed water quality parameters in the dry (red) and wet (blue) season along the Musi River.** The river can be divided into three stretches: (a) one sampling site upstream (light blue), (b) seven sampling sites within the city (dark blue) and (c) two sampling sites downstream of the city (medium blue).

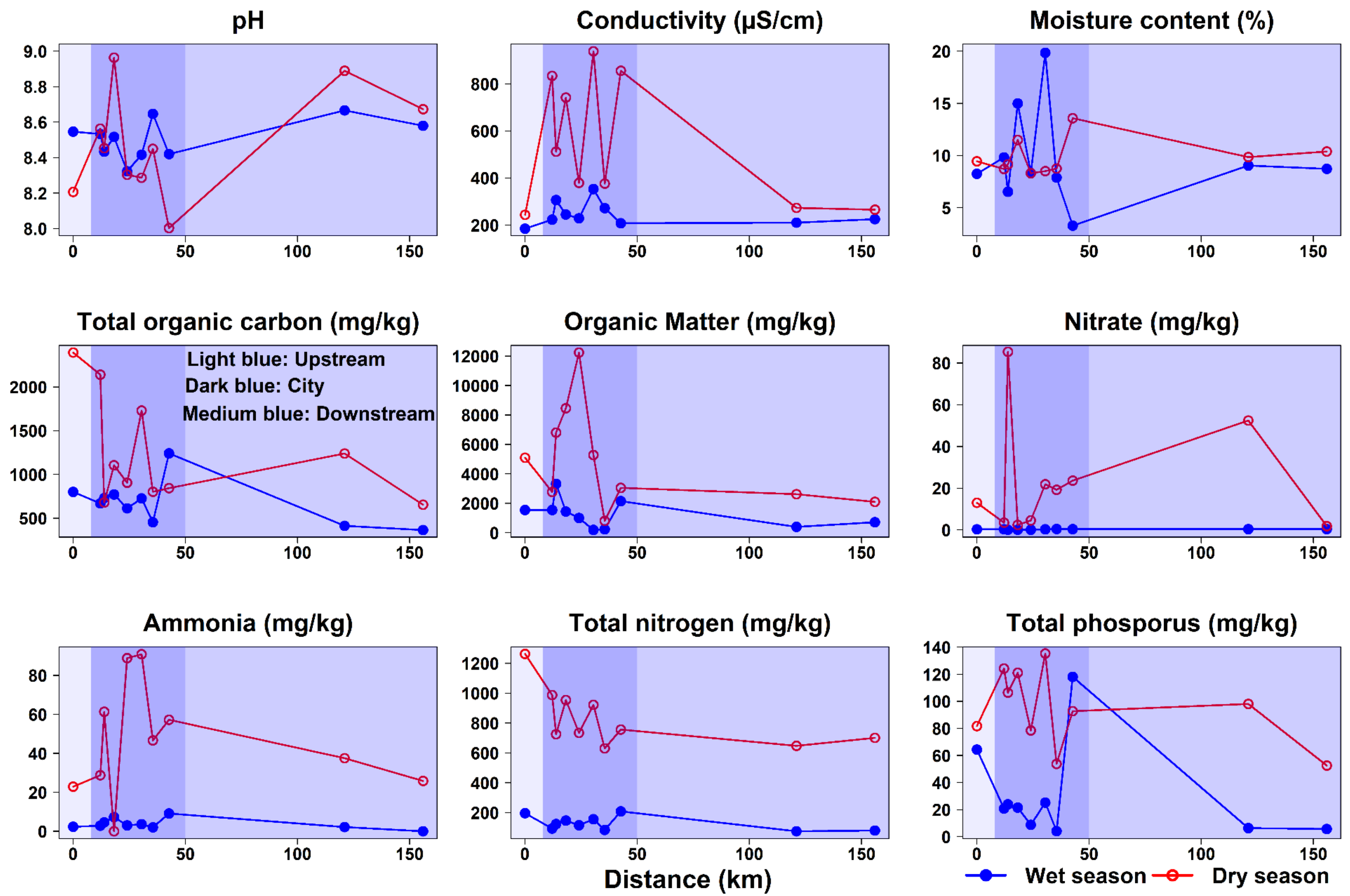


**Fig. S2. Sediment samples: All analysed sediment quality parameters in the dry (red) and wet (blue) season along the Musi River.** The river can be divided into three stretches: (a) one sampling site upstream (light blue), (b) seven sampling sites within the city (dark blue) and (c) two sampling sites downstream of the city (medium blue).

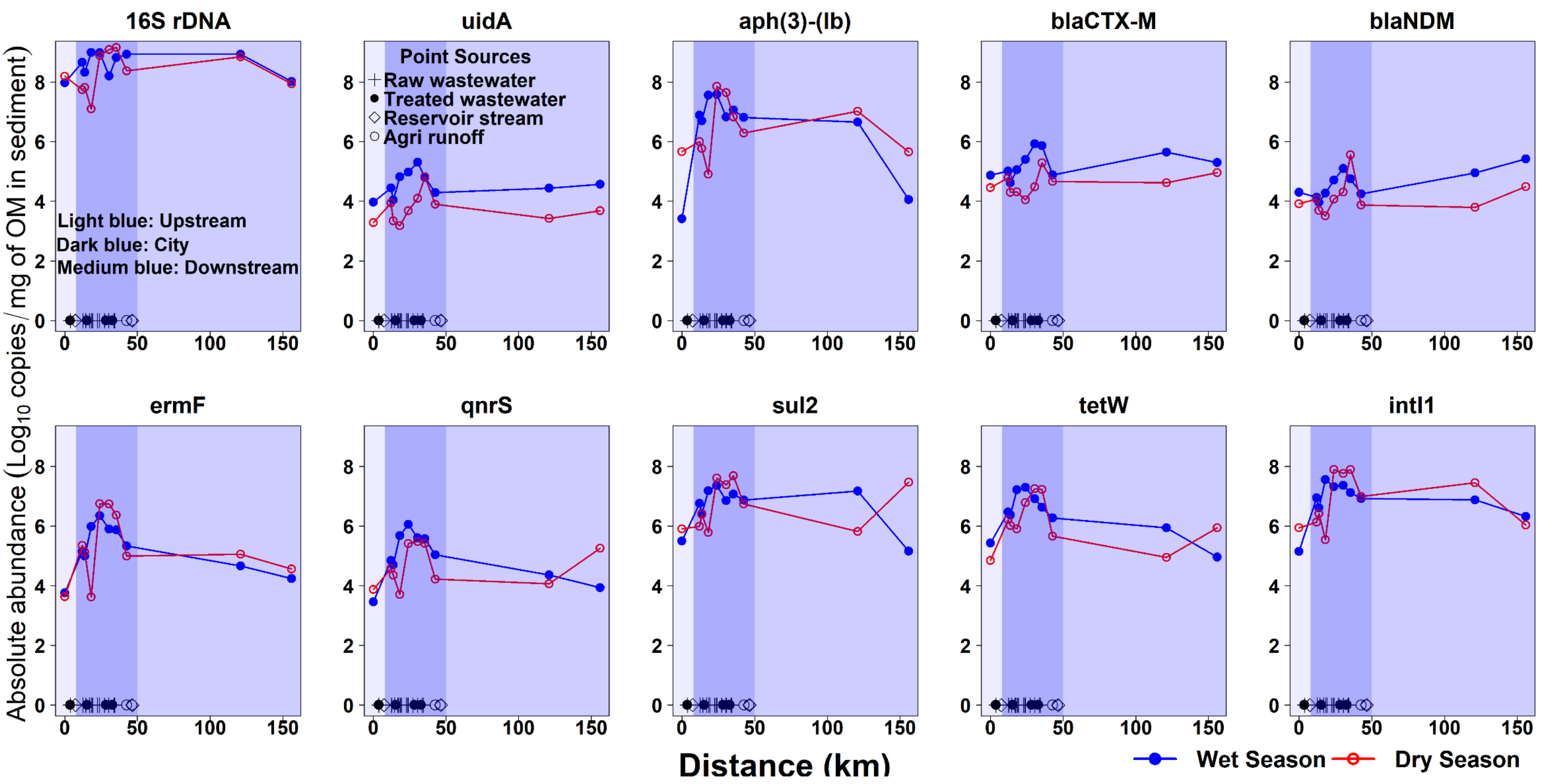


**Fig. S3. Absolute abundance of genes in sediment samples along the Musi River in the dry (red) and wet (blue) seasons.** All panels have the same scale showing log transformed absolute abundances of 16S rDNA, *uidA*, *intI1* and ARGs in the sediment column of the Musi River. The river can be divided into three stretches: (a) one sampling site upstream (light blue), (b) seven sampling sites within the city (dark blue) and (c) two sampling sites downstream of the city (medium blue). Point-sources of raw wastewater, treated wastewater, reservoir flows, and agricultural runoff are indicated by symbols.

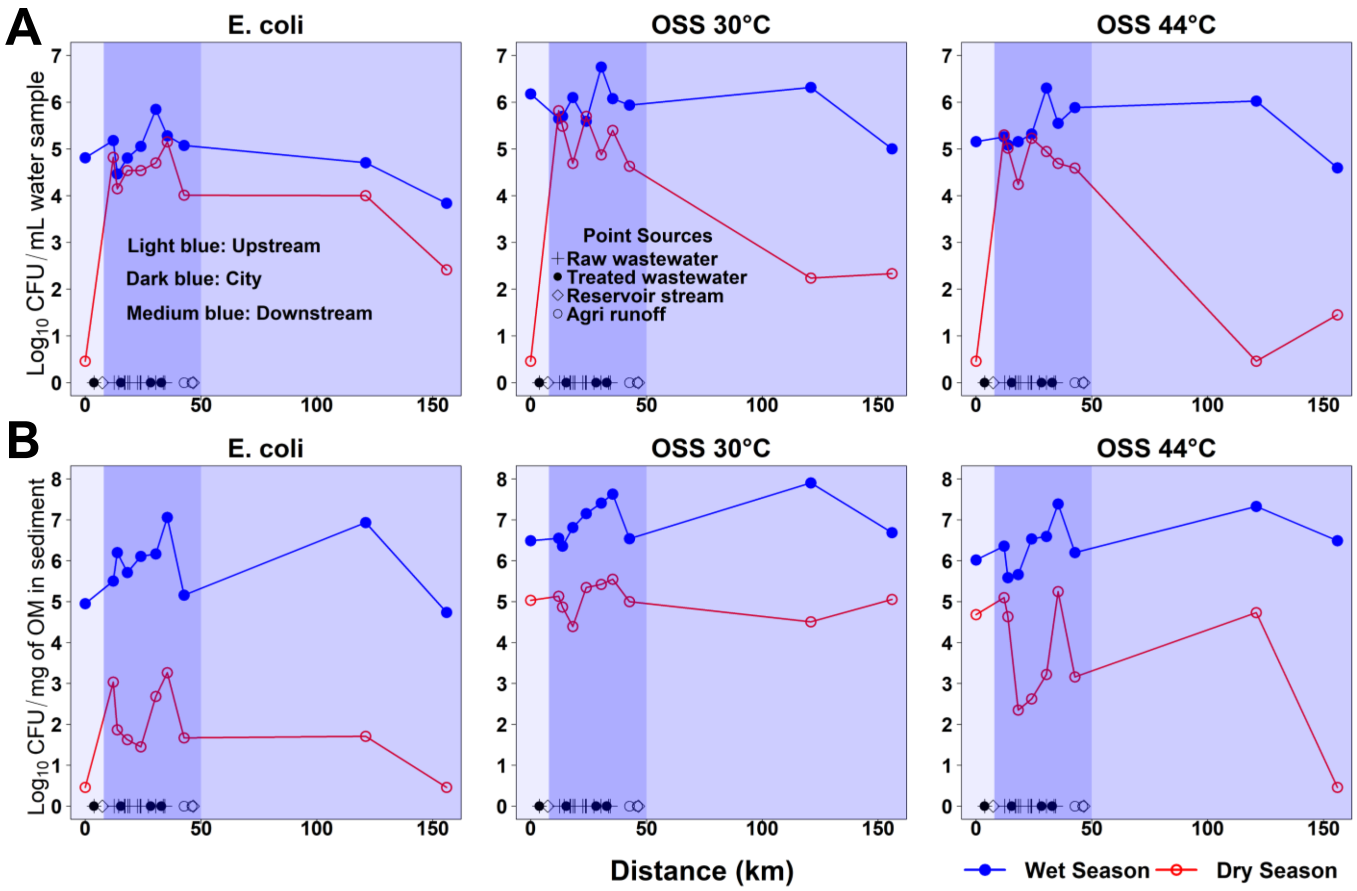


**Fig. S4. Viable counts (absolute abundances) of *E. coli* (blue colonies on TBX at 44°C) and aerobic heterotrophic bacteria on OECD synthetic sewage (OSS) agar at different temperatures (30°C and 44°C) in water (A) and sediment (B) along the river in the dry (red) and wet seasons (blue).** The river can be divided into three stretches: (a) one sampling site upstream (light blue), (b) seven sampling sites within the city (dark blue) and (c) two sampling sites downstream of the city (medium blue). Point-sources of raw wastewater, treated wastewater, reservoir flows, and agricultural runoff are indicated by symbols.

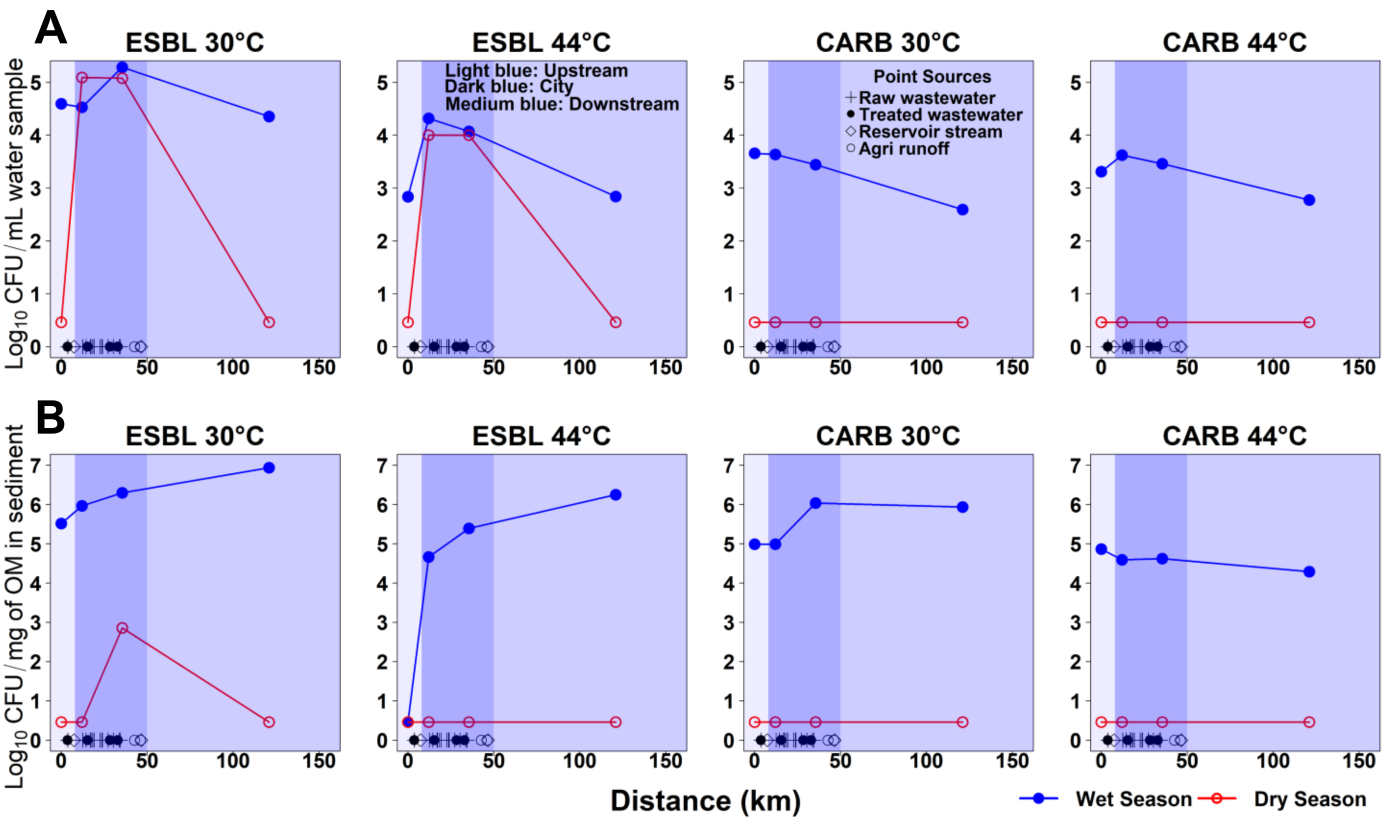


**Fig. S5. Viable counts (absolute abundances) of antibiotic-resistant aerobic heterotrophic bacteria on OECD synthetic sewage (OSS) agar supplemented with Extended Spectrum β-lactams (ESBL) and Carbapenems (CARB) at different temperatures (30°C and 44°C) in water (A) and sediment (B) along the river in the dry (red) and wet seasons (blue).** For ARB analysis, we selected four of our 10 sites: upstream (1); city (2 and 7); and downstream (9). The river can be divided into three stretches: (a) upstream (light blue), (b) within the city (dark blue) and (c) downstream of the city (medium blue). Point-sources of raw wastewater, treated wastewater, reservoir flows, and agricultural runoff are indicated by symbols.

### *1.1. Bacterial plate counts and their correlations with genes*

Comparing counts with different media and temperatures to the highest counts on OSS medium at 30°C as a reference, counts on TBX agar at 44°C and ESBL agar at 30°C were roughly 1 log10 lower and counts on CARB agar at 30°C were roughly 2 log10 lower (**Fig. S6 A-B**). Comparing counts at the higher temperature of 44°C to counts on the same medium at 30°C, they were roughly 0.5 log10 lower for OSS, 1 log10 lower for ESBL and only 0.25 log10 lower for CARB, suggesting that only about one tenth of ESBL bacteria but about half of the carbapenem resistant bacteria grow at 44°C (**Fig. S6 C-D**).

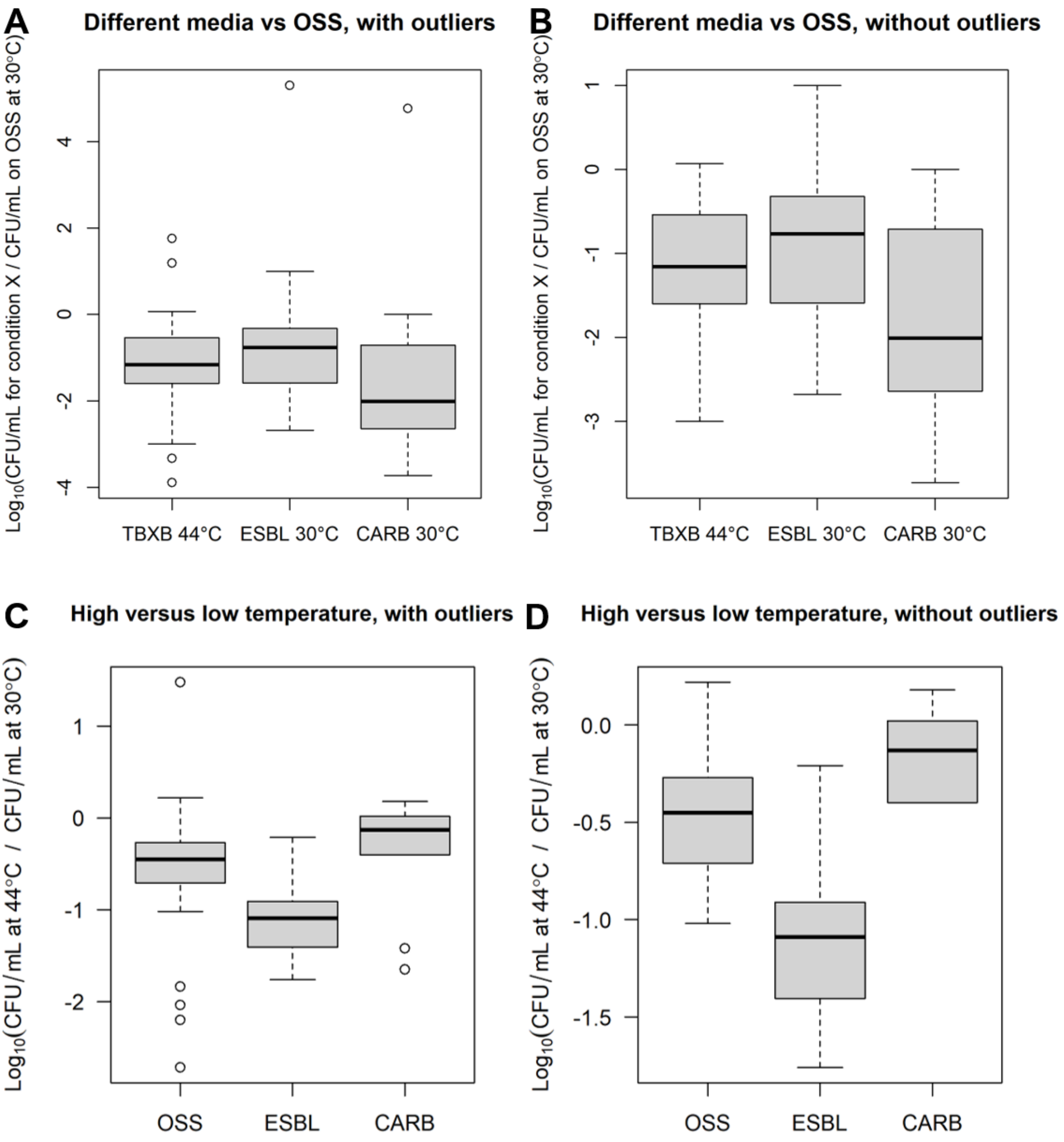


**Fig. S6. Comparison of viable counts with different media and temperatures in terms of log(ratio). (A-B)** Viable counts on the more selective media were lower than on OSS at 30°C. **(C-D)** Viable counts at 44°C were lower than at 30°C on the same medium. Acronyms: OSS: OECD Synthetic Sewage medium; TBXB: Tryptone Bile X-glucuronide blue colonies; ESBL: Extended Spectrum β-lactamase producing bacteria; CARB: Carbapenemase producing bacteria.

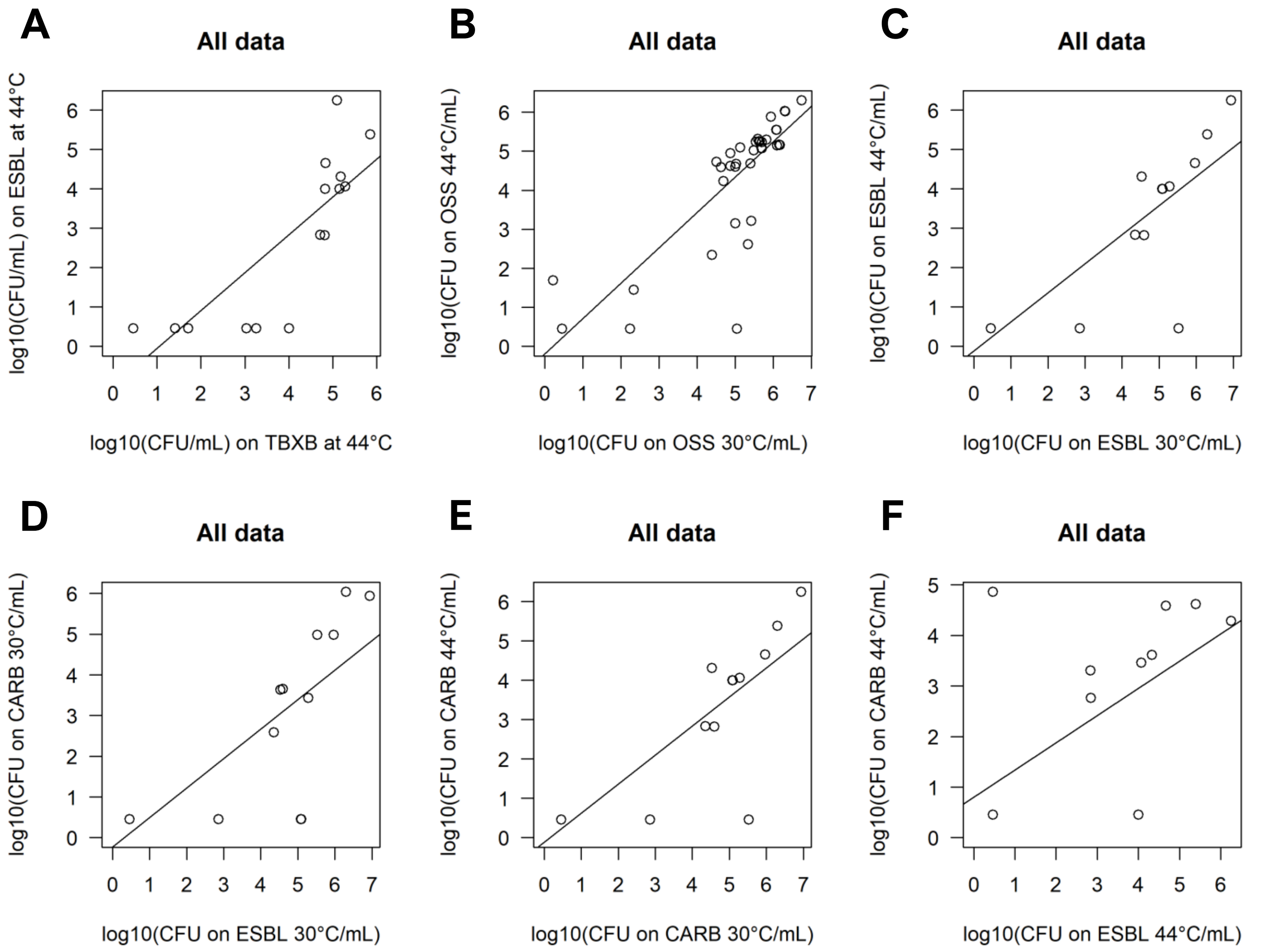


**Fig. S7. Effect of medium and temperature on viable counts.** Linear regressions of log-transformed viable counts using combined water and sediment data. Acronyms: OSS: OECD Synthetic Sewage medium; TBXB: Tryptone Bile X-glucuronide blue colonies; ESBL: Extended Spectrum β-lactamase producing bacteria; CARB: Carbapenemase producing bacteria. Statistics are given in **Table S1**.

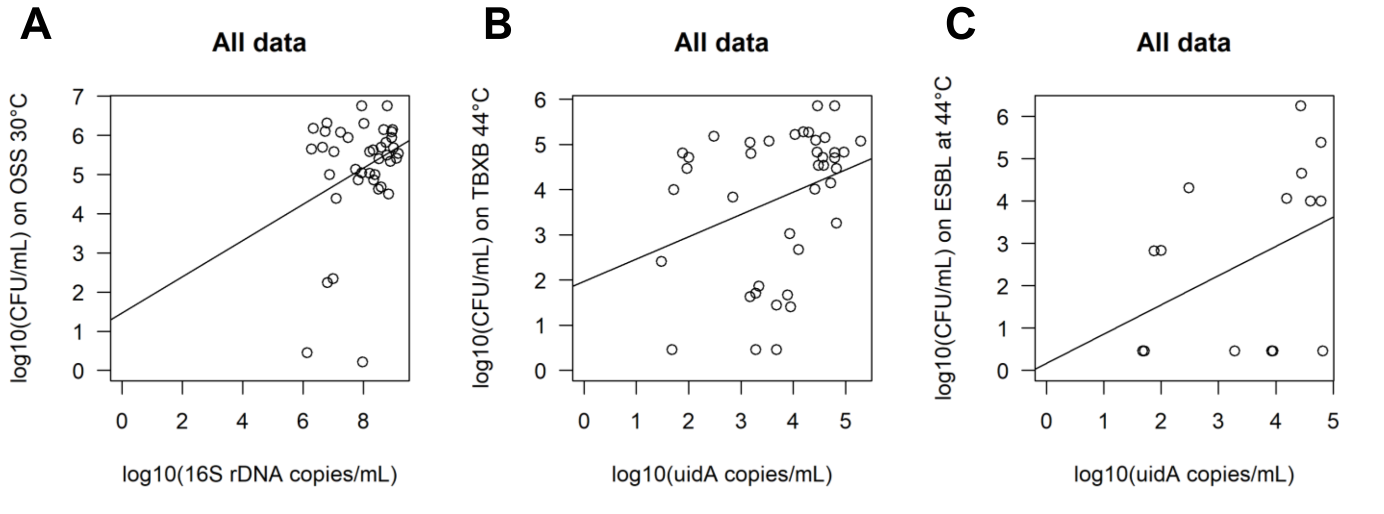


**Fig. S8. Linear regressions of log-transformed viable counts with different media and temperatures versus gene copies (16S rDNA and *uidA*)** using combined water and sediment data. Acronyms: see **Fig. S7**. Statistics are given in **Table S1**.

**Table S1. Linear regression analysis of (a) log-transformed viable counts for various media and temperature conditions (Fig. S7) and (b) log-transformed viable counts versus copies of 16S rDNA or *uidA* genes (Fig. S8).** Water and sediment data combined (all data). If the response variable is a fraction of the explanatory variable for untransformed data, we are expecting a slope of 1 and a negative y-intercept corresponding to the log of the fraction for the log transformed data.

| Linear Regression Analysis | Intercept | Intercept SE | Intercept *p*-value | Slope | Slope SE | Slope *p*-value | Adjusted $R^2$ |
|---|---|---|---|---|---|---|---|
| **(a)** TBXB 44°C vs ESBL 44°C | -1.0097 | 0.7287 | 0.19 | 0.96 | 0.1760 | 8.35E-05 | 0.6579 |
| OSS 44°C vs OSS 30°C | -0.1918 | 0.5715 | 0.74 | 0.90 | 0.1071 | 2.99E-10 | 0.6433 |
| ESBL 44°C vs ESBL 44°C | -0.1141 | 0.5447 | 0.84 | 0.74 | 0.1251 | 3.88E-05 | 0.6926 |
| CARB 44°C vs CARB 44°C | 0.1837 | 0.1432 | 0.22 | 0.83 | 0.0441 | 2.55E-11 | 0.9590 |
| ESBL 30°C vs CARB 30°C | -0.2275 | 0.6572 | 0.73 | 0.72 | 0.1510 | 2.83E-04 | 0.5950 |
| ESBL 44°C vs CARB 44°C | 0.8034 | 0.6233 | 0.22 | 0.54 | 0.1887 | 1.29E-02 | 0.3216 |
| **(b)** 16S rDNA vs OSS 30°C | 3.7707 | 1.4220 | 0.012 | 0.20 | 0.1768 | 0.2577 | 0.0086 |
| *uidA* vs TBXB 44°C | 2.2966 | 0.8223 | 0.0083 | 0.4527 | 0.2107 | 0.0384 | 0.0891 |
| *uidA* vs ESBL 44°C | 0.0308 | 0.9658 | 0.9752 | 1.0524 | 0.2674 | 0.0034 | 0.5916 |

### *1.2. Stretches and seasons were significantly different*

Here we provide an analysis of the PERMANOVA results in **Table S2** and how they can inform the LDA analysis in section 2.4 in the main text. Further results are in Supplementary Data 2.

#### *Water samples*

For genes, Stretch was always significant, for all individual genes and ARGs as a unit. Likewise for Season, apart from *tetW*. The Interaction Stretch:Season was significant for ARGs as a unit and most genes but not *aph(3")-Ib*, $bla_{\mathrm{NDM}}$ and *tetW*. Stretch, Season and Interaction together explained >0.7 of the variation, from 0.73 to 0.88 (adjusted $R^2$ values). Stretch explained more than Season, and Season explained a similar extent of variation as Interaction.

For water quality parameters, stretch was significant apart from nitrite, suggesting that nitrite was not informative, consistent with LDA results that nitrite can be removed without affecting the discrimination. Season was significant apart from DO and nitrite, suggesting DO differs less between seasons but strongly between stretches, which makes it valuable for discriminating stretches. The Interaction was significant apart from COD and nitrite. Stretch explained more of the variation than Season for most variables, especially for DO, confirming the above. The exception was ammonium (but not TN), where variation was much better explained by season. The interaction in some cases explained as much of the variation as the leading factor (pH) or the secondary factor (nitrate and ammonium). The sum of $R^2$ was highest for ammonium, DO and TN (>0.9) and lowest for nitrite and COD (0.5-0.6). This suggests that ammonium and DO and TN had the strongest signal, with DO for Stretch and ammonium for Season and TN in between.

For viable counts, Stretch, Season and Interaction were all significant for all counts that could be analysed (not the ARBs, for which not enough locations were sampled). The sum of the adjusted $R^2$ values was >0.86, with Stretch and Season and Interaction explaining similar proportions of the variance. Counts of *E. coli* (blue colonies on TBX agar at 44°C) were more Stretch than Season dependent, while for counts on OSS at 30°C and at 44°C, Stretch and Season were similarly important.

***Sediment samples***

In the sediment, the pattern was generally more complex. For genes, the 16S gene was neither significantly different between Stretches nor Seasons and the sum of the adjusted $R^2$ values was the lowest of all at 0.14, suggesting a fairly uniform presence of 16S as expected. For ARGs as a unit and for the individual ARGs *intI1, aph(3")-Ib, ermF, qnrS* and *tetW*, Stretch was significant, but Season and Interaction were not. The opposite (Stretch insignificant but Season significant) was found for *uidA* and $bla_{\text{CTX-M}}$, while none were significant for 16S, $bla_{\text{NDM}}$ and *sul2*. The sums of the adjusted $R^2$ values were 0.14-0.61, so a lot of variation was unexplained. Stretch explained >0.1 of the variation seven times, Season three times and Interaction two times.

For sediment quality parameters, Stretch was mostly insignificant, apart from Conductivity and TN. Season on the other hand was generally significant, with pH the only exception. The Interaction was generally not significant, with TN the only exception. TN was always significant. The sum of the adjusted $R^2$ values was highest for TN (0.96) and low for nitrate and pH (~0.3), intermediate for the rest (0.59-0.69). Adjusted $R^2$ values for Season were generally higher than for Stretch, apart from pH. Adjusted $R^2$ values for Interactions were low.

For viable counts, there were not enough data to analyse ARBs. Stretch was significant for counts on antibiotic free media apart from OSS44. Season and Interaction were always significant. The sums of the adjusted $R^2$ values were high (> 0.88), apart from OSS44 at 0.6. Adjusted $R^2$ values for Season were higher than for Stretch, apart from OSS30, where Stretch and Interaction were higher.

**Table S2. Results of PERMANOVA analysis for the individual and aggregate variables ARGs, water quality (WQ), sediment quality (SQ), and viable counts (CFUs).** These values are based on Euclidean distance, for Bray-Curtis dissimilarity base values see Supplementary Data 2.

| Sets | Water | Stretch | | Season | | Interaction | | Sum | Sediment | Stretch | | Season | | Interaction | | Sum |
|---|---|---|---|---|---|---|---|---|---|---|---|---|---|---|---|---|
| | | $R^2$ | *P value* | $R^2$ | *P value* | $R^2$ | *P value* | $R^2$ | | $R^2$ | *P value* | $R^2$ | *P value* | $R^2$ | *P value* | $R^2$ |
| **Genes** | 16S rDNA | 0.378 | 0.0001 | 0.343 | 0.0001 | 0.155 | 0.0010 | 0.876 | 16S rDNA | 0.052 | 0.683 | 0.057 | 0.342 | 0.034 | 0.768 | 0.143 |
| | *uidA* | 0.587 | 0.0001 | 0.096 | 0.0158 | 0.141 | 0.0097 | 0.824 | *uidA* | 0.112 | 0.180 | 0.479 | 0.003 | 0.006 | 0.905 | 0.598 |
| | *aph3-Ib* | 0.715 | 0.0001 | 0.085 | 0.0068 | 0.073 | 0.0524 | 0.872 | *aph3-Ib* | 0.377 | 0.023 | 0.000 | 0.980 | 0.179 | 0.108 | 0.556 |
| | *bla*$_{\text{CTX-M}}$ | 0.486 | 0.0002 | 0.188 | 0.0031 | 0.135 | 0.0136 | 0.809 | *bla*$_{\text{CTX-M}}$ | 0.060 | 0.461 | 0.426 | 0.006 | 0.007 | 0.915 | 0.493 |
| | *bla*$_{\text{NDM}}$ | 0.551 | 0.0001 | 0.151 | 0.0093 | 0.079 | 0.0936 | 0.782 | *bla*$_{\text{NDM}}$ | 0.089 | 0.418 | 0.169 | 0.085 | 0.073 | 0.483 | 0.331 |
| | *ermF* | 0.454 | 0.0003 | 0.209 | 0.0036 | 0.120 | 0.0364 | 0.784 | *ermF* | 0.458 | 0.007 | 0.000 | 0.996 | 0.009 | 0.900 | 0.467 |
| | *qnrS* | 0.546 | 0.0002 | 0.123 | 0.0116 | 0.126 | 0.0210 | 0.795 | *qnrS* | 0.369 | 0.011 | 0.037 | 0.313 | 0.114 | 0.219 | 0.520 |
| | *sul2* | 0.673 | 0.0001 | 0.091 | 0.0090 | 0.099 | 0.0239 | 0.863 | *sul2* | 0.251 | 0.114 | 0.001 | 0.892 | 0.033 | 0.728 | 0.285 |
| | *tetW* | 0.549 | 0.0002 | 0.067 | 0.0784 | 0.112 | 0.0671 | 0.728 | *tetW* | 0.573 | 0.001 | 0.030 | 0.319 | 0.011 | 0.832 | 0.614 |
| | *intI1* | 0.662 | 0.0002 | 0.077 | 0.0195 | 0.112 | 0.0181 | 0.851 | *intI1* | 0.344 | 0.043 | 0.000 | 0.967 | 0.037 | 0.669 | 0.381 |
| | **All ARGs** | **0.577** | **0.0001** | **0.129** | **0.0085** | **0.104** | **0.0307** | 0.809 | **All ARGs** | **0.364** | **0.006** | **0.045** | **0.277** | **0.080** | **0.358** | 0.489 |
| **WQ/SQ** | pH | 0.356 | 0.0004 | 0.173 | 0.0003 | 0.341 | 0.0023 | 0.870 | pH | 0.244 | 0.1187 | 0.005 | 0.7491 | 0.090 | 0.4159 | 0.338 |
| | DO | 0.897 | 0.0001 | 0.003 | 0.4001 | 0.053 | 0.0049 | 0.953 | Conductivity | 0.195 | 0.0446 | 0.386 | 0.0002 | 0.111 | 0.1213 | 0.692 |
| | COD | 0.280 | 0.0401 | 0.223 | 0.0144 | 0.063 | 0.3789 | 0.566 | TOC | 0.202 | 0.0610 | 0.285 | 0.0047 | 0.104 | 0.2282 | 0.591 |
| | Nitrate | 0.398 | 0.0136 | 0.192 | 0.0014 | 0.244 | 0.0422 | 0.834 | Nitrate | 0.008 | 0.9483 | 0.281 | 0.0275 | 0.007 | 0.8660 | 0.296 |
| | Ammonium | 0.140 | 0.0001 | 0.674 | 0.0001 | 0.148 | 0.0002 | 0.962 | Ammonium | 0.050 | 0.4021 | 0.537 | 0.0005 | 0.031 | 0.5667 | 0.618 |
| | Nitrite | 0.335 | 0.1133 | 0.076 | 0.1380 | 0.109 | 0.2782 | 0.520 | | | | | | | | |
| | | | | | | | | | TP | 0.060 | 0.3841 | 0.527 | 0.0014 | 0.031 | 0.5967 | 0.619 |
| | TN | 0.492 | 0.0001 | 0.253 | 0.0001 | 0.154 | 0.0008 | 0.898 | TN | 0.059 | 0.0014 | 0.869 | 0.0001 | 0.028 | 0.0212 | 0.956 |
| | **All WQ** | **0.414** | **0.0001** | **0.228** | **0.0001** | **0.159** | **0.0004** | 0.800 | **All SQ** | **0.117** | **0.0803** | **0.413** | **0.0001** | **0.057** | **0.4741** | 0.587 |
| **Counts** | *E. coli* | 0.409 | 0.0049 | 0.205 | 0.0002 | 0.246 | 0.0183 | 0.859 | *E. coli* | 0.214 | 0.0001 | 0.656 | 0.0001 | 0.044 | 0.0369 | 0.914 |
| | OSS 30°C | 0.311 | 0.0012 | 0.327 | 0.0001 | 0.294 | 0.0007 | 0.931 | OSS 30°C | 0.453 | 0.0003 | 0.027 | 0.0344 | 0.457 | 0.0005 | 0.936 |
| | OSS 44°C | 0.360 | 0.0013 | 0.294 | 0.0001 | 0.282 | 0.0040 | 0.936 | OSS 44°C | 0.073 | 0.2931 | 0.245 | 0.0098 | 0.285 | 0.0253 | 0.603 |
| | **All CFU** | **0.272** | **0.0001** | **0.338** | **0.0001** | **0.325** | **0.0002** | 0.934 | **All CFU** | **0.094** | **0.0111** | **0.543** | **0.0001** | **0.238** | **0.0002** | 0.875 |

### *1.3. Estimates of municipal wastewater discharges through various drains into the Musi River*

To estimate municipal wastewater discharges into the Musi River, three different types of data were combined: (a) local population sizes from ward census data and population density maps, (b) drainage network maps, and (c) wastewater production rates. There were no wastewater drains or WWTP effluents upstream or downstream of the city.

**(a) Ward census data and population density maps:** The population density ward map of Hyderabad city was obtained from the Greater Hyderabad Municipal Corporation, dividing the city into 150 wards, each identified as a significant contributor to wastewater discharge into the Musi River. The population census data was sourced from the Census of India (MoHA 2022, https://censusindia.gov.in/census.website/data/census-tables) and supplemented with information from the 'datameet Google group' due to changes in ward names post 2014 (https://groups.google.com/g/datameet/c/DiE3Wtyup0E). The projected population data for 2022 were acquired from the World Population Review (https://worldpopulationreview.com/world-cities/hyderabad-population). Additionally, WorldPop provided a population density map for areas outside Hyderabad's administrative boundaries (https://data.humdata.org/dataset/worldpop-population-density-for-india).

**(b) Drainage network of the Musi River:** The drainage network was derived from a 30-metre Shuttle Radar Topography Mission (SRTM) Digital Elevation Model (DEM) and manually traced using Google Earth. The DEM primarily identified non-visible drainage paths, which were then validated manually. The combined network, illustrated in **Fig. S13**, was deemed suitable for discharge calculations.

**(c) Wastewater production rates:** Wastewater production per capita per day (in litre per capita per day, lcpd) for Hyderabad was estimated from all data on population size, water supply and wastewater discharge rates we were able to find on government websites, publications, research organisation reports and newspapers from 2001 to 2022 (see **Supplementary Data 4**). The direct estimation method was to divide the total wastewater production rate in the city by the city’s population size for the corresponding year (interpolated if necessary). The indirect estimation method was to first convert drinking water supply rates for the city into wastewater production rates, assuming that 80% of drinking water ends up as sewage and then divide this by population size as for the direct method. The two methods were reasonably consistent, so data were aggregated (averages and standard errors calculated for years with several data points) and a linear regression of wastewater production versus year was used to estimate wastewater production in 2022 (**Fig. S14, Supplementary Data 4**). The average sewage generation in 2022 was 150 ± 25 lpcd and 150 lpcd was used for further calculations. The following formula was used to calculate sewage discharge for a particular drain based on the population size from (a) in the catchment feeding into the drain identified from the topography in (b):

$$\text{Wastewater discharge (cumecs)} = \frac{150\ \text{lpcd} \times \text{corresponding ward population}}{24 \times 3600 \times 1000} \quad \text{(S1)}$$

Where lcpd denotes litre per capita per day and 24×3600×1000 converts litres per day into cubic metres per second (cumecs).

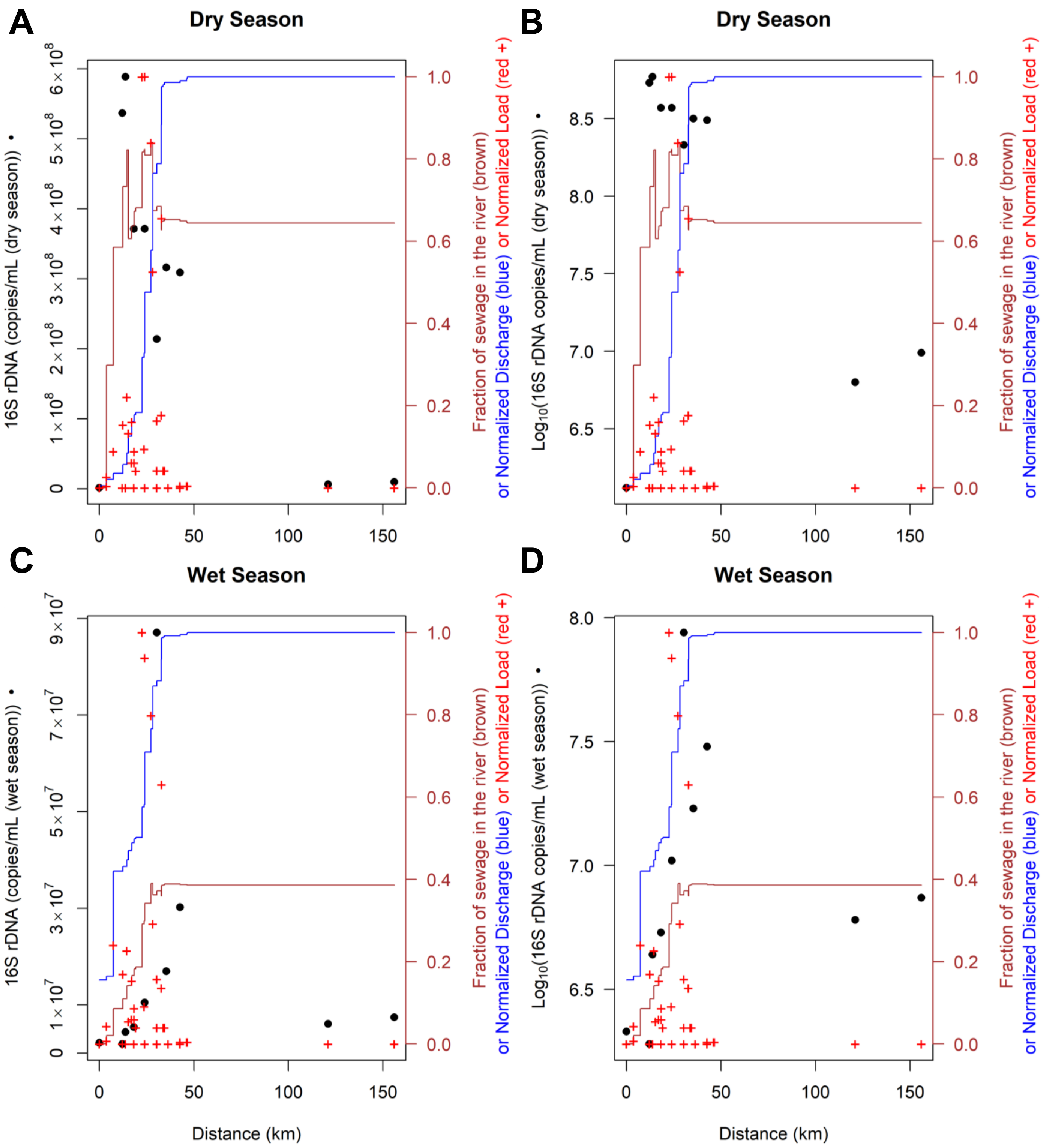


**Fig. S9. Direct comparison of the absolute abundance of the 16S gene in the river with the estimated wastewater derived fraction in both dry and wet seasons.** The hypothetical tracer concentration was set to 1 in raw wastewater and to 0.16 in treated wastewater to track wastewater loads. The tracer was assumed to be inert in contrast to 16S rDNA levels (•), which declined along the river. Normalized discharge (blue stairs), normalized loadings (+) and the fraction of wastewater in the river based on tracer mass balance calculations (brown stairs). **(A, C)** Comparing the actual concentration of the 16S gene (not log-transformed) with the wastewater fraction being the assumed main source of the 16S gene-carrying bacteria. **(B, D)** Comparing $Log_{10}$ transformed 16S concentrations with the tracer concentrations as other figures are showing log-transformed data, visualising the relationship between untransformed and transformed data.

**Table S3. Antibiotic concentrations (µg/L) in the water of the Musi River observed in other studies. All sampling was done in the dry season.** The study by Wilkinson et al.[1] measured 13 antibiotics at 18 sites in Hyderabad but did not report locations and date of sampling so was not included in this table. They detected three out of the 13 antibiotics in 6-7 locations, with average concentrations of 0.30 for ciprofloxacin, 0.19 for sulfamethoxazole and 0.020 for trimethoprim, which are similar to the range of values reported in the studies of Konda et al.[2,3] for city samples.

| Reference | Sampling site in Musi River | Sampling Date | Stretch | Latitude | Longitude | Ciprofloxacin | Enrofloxacin | Norfloxacin | Difloxacin | Pefloxacin | Lomefloxacin | Ofloxacin | Levofloxacin | Sulfamethoxazole | Sulphapyridine | Clarithromycin | Trimethoprim |
|---|---|---|---|---|---|---|---|---|---|---|---|---|---|---|---|---|---|
| (Gothwal and Shashidhar, 2017)[4] | Osman Sagar | Apr-2015 | Upstream | 17°22′50.16″ N | 78°18′59.4″ E | 27.31 | 3.77 | 26.68 | 3.11 | 2.36 | 5.17 | 3.72 | | | | | |
| | Langar Hauz, Bapu Ghat | | City | 17°22′20.24″ N | 78°24′45.72″ E | 43.76 | 12.46 | 58.91 | 8.37 | 3.73 | 4.57 | 58.47 | | | | | |
| | Nayapul | | City | 17°22′4.79″ N | 78°27′25.19″ E | 245.55 | 14.78 | 112.9 | 8.73 | 3.279 | 9.34 | 210.4 | | | | | |
| | Musaram Baug | | City | 17°22′47.74″ N | 78°31′2.28″ E | 5,528.90 | 123.40 | 217.50 | 44.34 | 37.74 | 7.80 | 318.10 | | | | | |
| | Peerzadiguda | | City | 17°23′20.03″ N | 78°35′52.79″ E | 1,999.60 | 44.38 | 148.54 | 17.45 | 8.89 | 10.32 | 204.57 | | | | | |
| | Pratap Singaram | | City | 17°22′48.9″ N | 78°40′1.2″ E | 662.34 | 22.07 | 92.98 | 5.5 | 10.35 | 6.78 | 199.55 | | | | | |
| | Pillaipalli | | Downstream | 17°23′5.6″ N | 78°44′15.32″E | 231.9 | 13.73 | 79.37 | 8.01 | 3.64 | 3.93 | 112.6 | | | | | |
| | Rudravalli | | Downstream | 17°24′22.53″ N | 78°47′9.6″ E | 86.89 | 15.82 | 49.41 | 2.026 | 4.26 | 4.73 | 56.6 | | | | | |
| | Suryapet Musi Reservoir | | Downstream | 17°14′22.38″ N | 79°30′59.07″ E | 10.72 | 14.67 | 31.84 | 7.38 | 1.85 | 4.56 | 4.526 | | | | | |
| (Lübbert et al., 2017)[5] | Upstream of Amberpet STP | 20-Nov-2016 | City | 17°23′08″ N | 78°30′36″E | 40.1 | | | | | | | 12.8 | | | | |
| | Weir in the Musi River, Hyderabad City, downstream of Amberpet STP | | City | 17°23′17″ N | 78°35′53″ | 44.7 | | | | | | | 10 | 10.6 | | 27.7 | |

| Reference | Sampling site in Musi River | Sampling Date | Stretch | Latitude | Longitude | Ciprofloxacin | Enrofloxacin | Norfloxacin | Difloxacin | Pefloxacin | Lomefloxacin | Ofloxacin | Levofloxacin | Sulfamethoxazole | Sulphapyridine | Clarithromycin | Trimethoprim |
|---|---|---|---|---|---|---|---|---|---|---|---|---|---|---|---|---|---|
| (Konda et al., 2022)[3] | Osman Sagar | Nov-2018 | Upstream | 17°23′40.2 “N | 78°17′29.6″ E | | | | | | | ND | | ND | ND | | ND |
| | Nagole | | City | 17°23′00.8 ″ N | 78°33′32.0″ E | | | | | | | 0.402 | | 0.038 | 0.108 | | 0.044 |
| (Konda et al., 2024)[2] | Osman Sagar | Jan-2018 | Upstream | 17º22.376’ N | 78º25.827’E | ND | ND | ND | | | | ND | | | | | |
| | Gudimalkapur | | City | 17º22.376’ N | 78º25.827’E | ND | ND | ND | | | | ND | | | | | |
| | Salarjung Museum | | City | 17º22.342’ N | 78º28.557’E | 0.05 | ND | ND | | | | ND | | | | | |
| | Hussain Sagar confluence | | City | 17º23.221’ N | 78º30.384’E | 0.74 | 0.07 | 0.02 | | | | 0.2 | | | | | |
| | D/S Amberpet STP | | City | 17º22.888’ N | 78º33.493’E | 0.05 | ND | ND | | | | ND | | | | | |
| | Nalla Cheruvu STP | | City | 17º23.510’ N | 78º34.577’E | 0.67 | 0.05 | 1.19 | | | | 0.08 | | | | | |
| | Pratap Singaram | | City | 17º22.825’ N | 78º40.039’E | 0.41 | ND | 0.33 | | | | ND | | | | | |

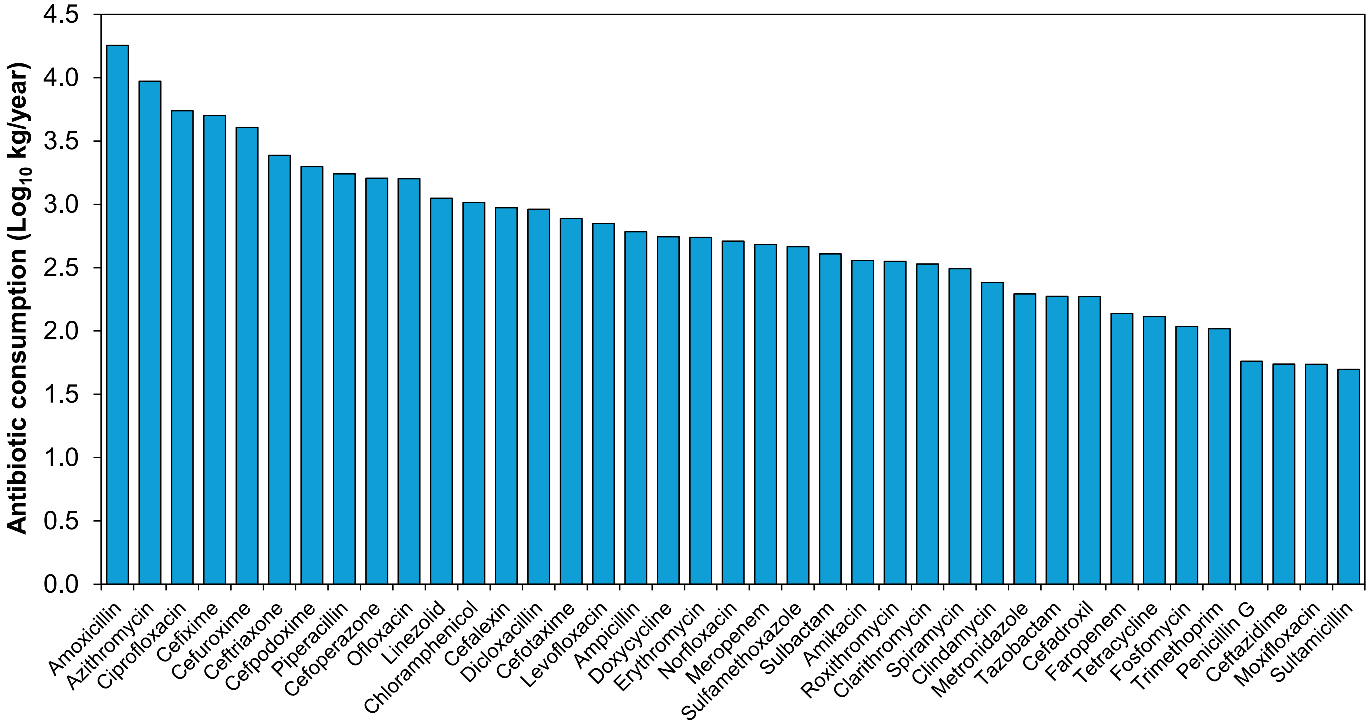


**Fig. S10. Ranked total consumption of antibiotics in Hyderabad in 2022, showing the top 39.** Data was calculated from sales data provided by IQVIA, assuming that sold antibiotics will be consumed. The population size of the Hyderabad metropolitan area in 2022 was about 10.5 million.

### *1.4. Patterns of environmental conditions*

Fifteen water quality parameters were measured in the Musi River, with a subset assessed in sediment samples (**Figs. S1-S2**). Before LDA, standardisation was needed, as boxplots displayed differences in scale and distribution across stretches and seasons (**Fig. S11**) and results of statistical testing for homogeneity of multivariate dispersions (within-group covariance, betadisper function in the vegan package) showed lack of homogeneity before standardisation but not after. To reduce the number of parameters, temperature was removed because it does not generalise to other regions, and conductivity, TDS, turbidity, TSS, VS, TP and TOC were removed due to collinearities.

The pattern of changes along the river for all water quality parameters is shown in **Fig. S1**. **Ammonia** in the dry season was low upstream, higher in the city and even higher downstream; this was in contrast to the 16S gene (**Fig. 1**), sewage pollution and AMR. In the wet season, ammonia was much lower and only slightly elevated in the city's second half, again different from the 16S gene. Thus, ammonia does not seem to be a suitable proxy for AMR pollution. **Nitrite** in the dry season had low values everywhere apart from one spike downstream (site 9), which had dropped down again at the last site, the Musi reservoir. In the wet season, nitrite had higher values, but not in the second half of the city. This suggests that low ammonia concentrations upstream and lack of oxygen in the city inhibited nitrification. Also, nitrite may be used in denitrification under low oxygen conditions. The higher oxygen levels in the wet season in the first half of the city allowed some nitrification. Nitrite may also be transported from further upstream in the wet season. **Nitrate** in the dry season was very similar to nitrite. In the wet season, it was also similar, apart from the upstream site Osman Sagar, which had much higher nitrate than nitrite concentrations. Consistent with the interpretation for nitrite, denitrification may have removed nitrite and nitrate in the city where oxygen was low, while the downstream increase may have been due to nitrification at higher oxygen levels. The drop at the last downstream site, the Musi reservoir, was possibly due to nitrate uptake by photosynthetic algae as longer residence time and larger surface area favour photosynthesis as discussed in the main paper. **TIN** (Total Inorganic Nitrogen, which is ammonia + nitrite + nitrate) was dominated by ammonia, so it showed the same pattern. None of the inorganic nitrogen compounds showed a pattern matching the 16S gene, suggesting they cannot be used as proxies for sewage and AMR pollution. **TN** (Total Nitrogen, measured with a TOC/TN analyser that includes all inorganic and organic nitrogen) showed a pattern that matched the 16S gene in both seasons, just dropping down a bit more downstream. This suggests that TN can be used as a proxy for sewage and AMR pollution. **TOC** (Total Organic Carbon) was similar to TN and the 16S gene but showed a noisier signal, so it gave a less clear distinction between upstream, city and downstream stretches. **TP** (Total Phosphate) showed a similar pattern to TOC in the dry season and also showed too much variation to be a good proxy. In the wet season, it had lower values and no clear pattern. **DO** (Dissolved Oxygen) showed a very clear pattern in the dry season: high upstream, plummeting in the city and gradually recovering downstream. In the wet season, the differences were less strong, and the decline in the city was more gradual, but the basic pattern was the same. DO was suitable as a proxy as it clearly discriminated upstream, city and downstream stretches in both seasons. **Conductivity** jumped up in the city but did not drop downstream, making it unsuitable for distinguishing the city from downstream sites that were already cleaner due to the river functioning as a treatment plant. For **pH**, the pattern inverted from the dry to the wet season; it was high upstream, dropped in the city, and went up again downstream during the dry season, while the wet season showed an opposite trend. **TDS** had the same pattern as conductivity (it is based on the same measurement). **Temperature** showed, as expected, no clear pattern along the river, while values were higher in the wet season. Since temperatures will vary between regions, it cannot be used as a general proxy for pollution. **Turbidity** was low upstream and downstream and high in the city; this pattern was clear in the dry season but less clear in the wet season.

Water quality parameters suggest nitrogen cycle changes along the Musi. The wastewater pollution from the city became decomposed by microbes, removing oxygen and releasing ammonia. With the drop in oxygen, denitrification kicked in and removed nitrite and nitrate in the city. Downstream, oxygen recovered, and nitrification set in while denitrification stopped, leading to a spike in nitrite and nitrate downstream of the city at site 9. These dropped again further downstream in the Musi reservoir (site 10), possibly due to the uptake of nitrate by photosynthetic algae as longer residence time and larger surface area favour photosynthesis (since 16S does not increase in the Musi reservoir, probably eukaryotic photosynthesis). Ammonia stays high due to the continuing decomposition of the still high organic carbon (TOC). Overall, the river functions as a sewage treatment plant for the city.

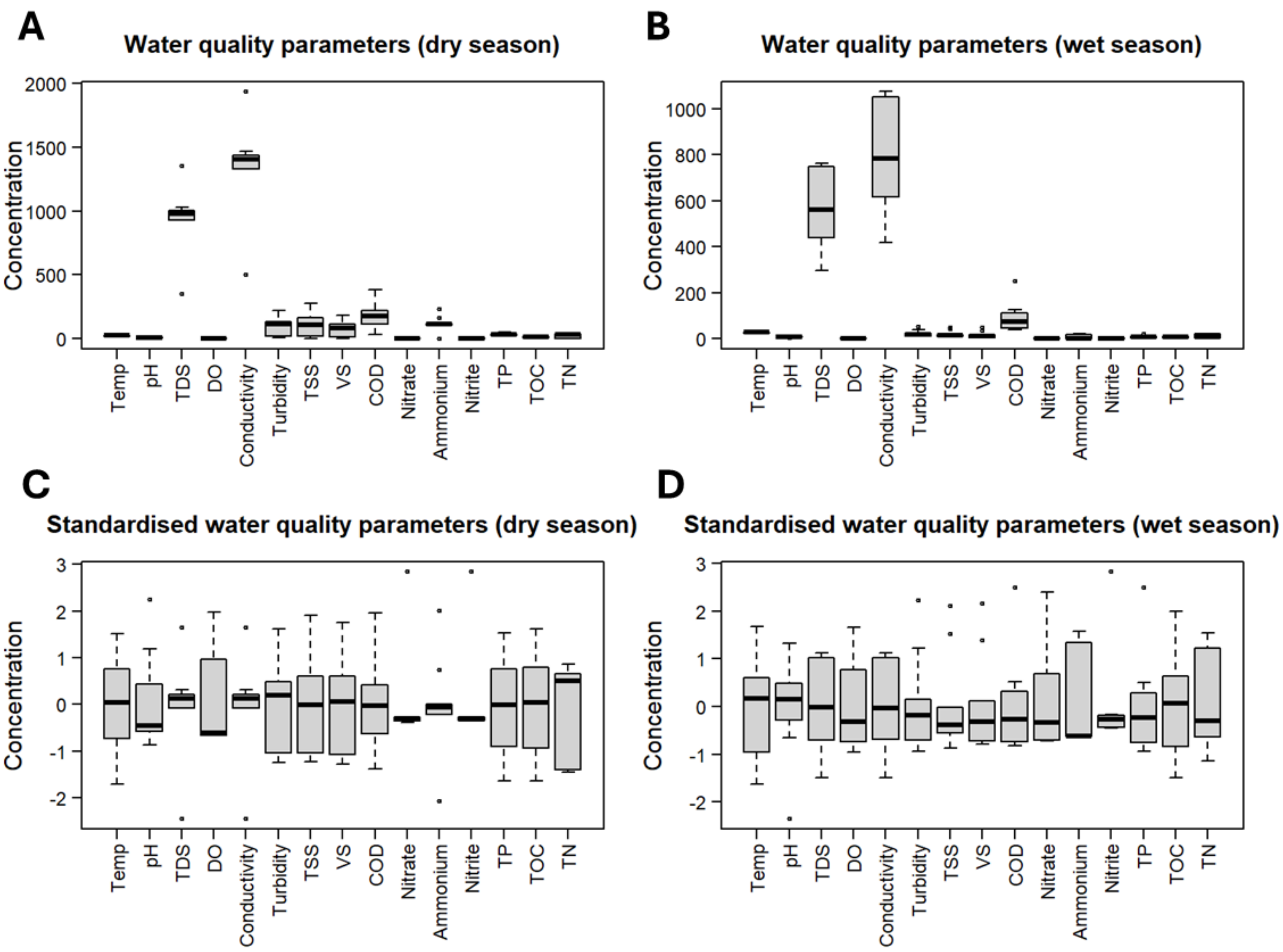


**Fig. S11. Boxplots of water quality parameters for both the dry (A, C) and wet (B, D) seasons of Musi River water samples. (A, B)** Boxplots of the untransformed and **(C, D)** standardised values of water quality parameters for both seasons. The boxes represent the 2nd and 3rd quartiles. The middle black line represents the median, and the whiskers represent the 1st and 4th quartiles. Black dots represent outlier values.

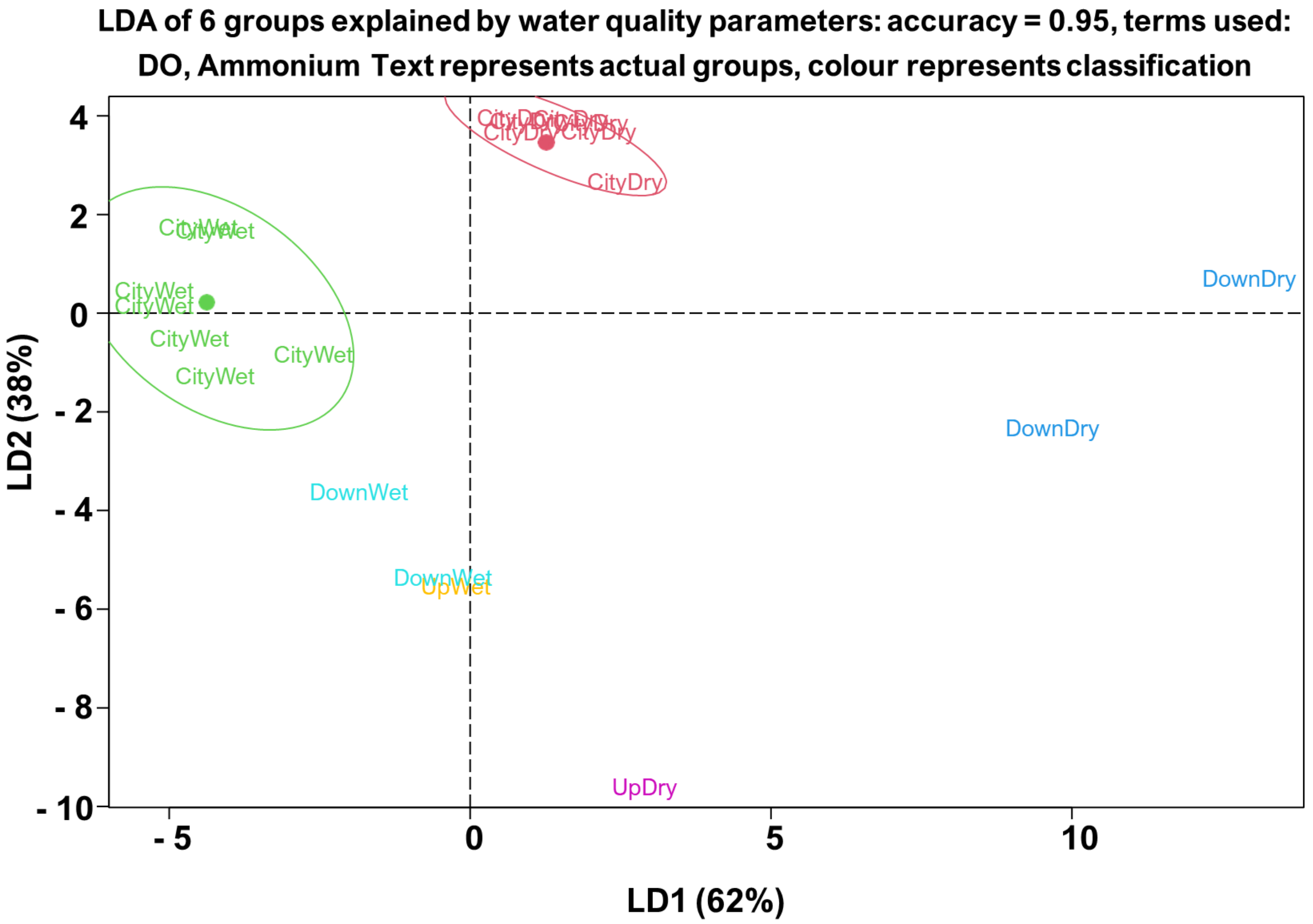


**Fig. S12. Linear discriminant analysis (LDA) showing that the three river stretches identified by clustering and ordination of ARGs can be discriminated by the two environmental characteristics DO and $NH_3$ in both seasons (resulting in 6 groups).** Text labels combine actual stretch (Up, City, Down) and season (Dry, Wet), while colours indicate the LDA classifications of the sites, with an accuracy of the classification of 95%. Ellipses show 95% confidence intervals.

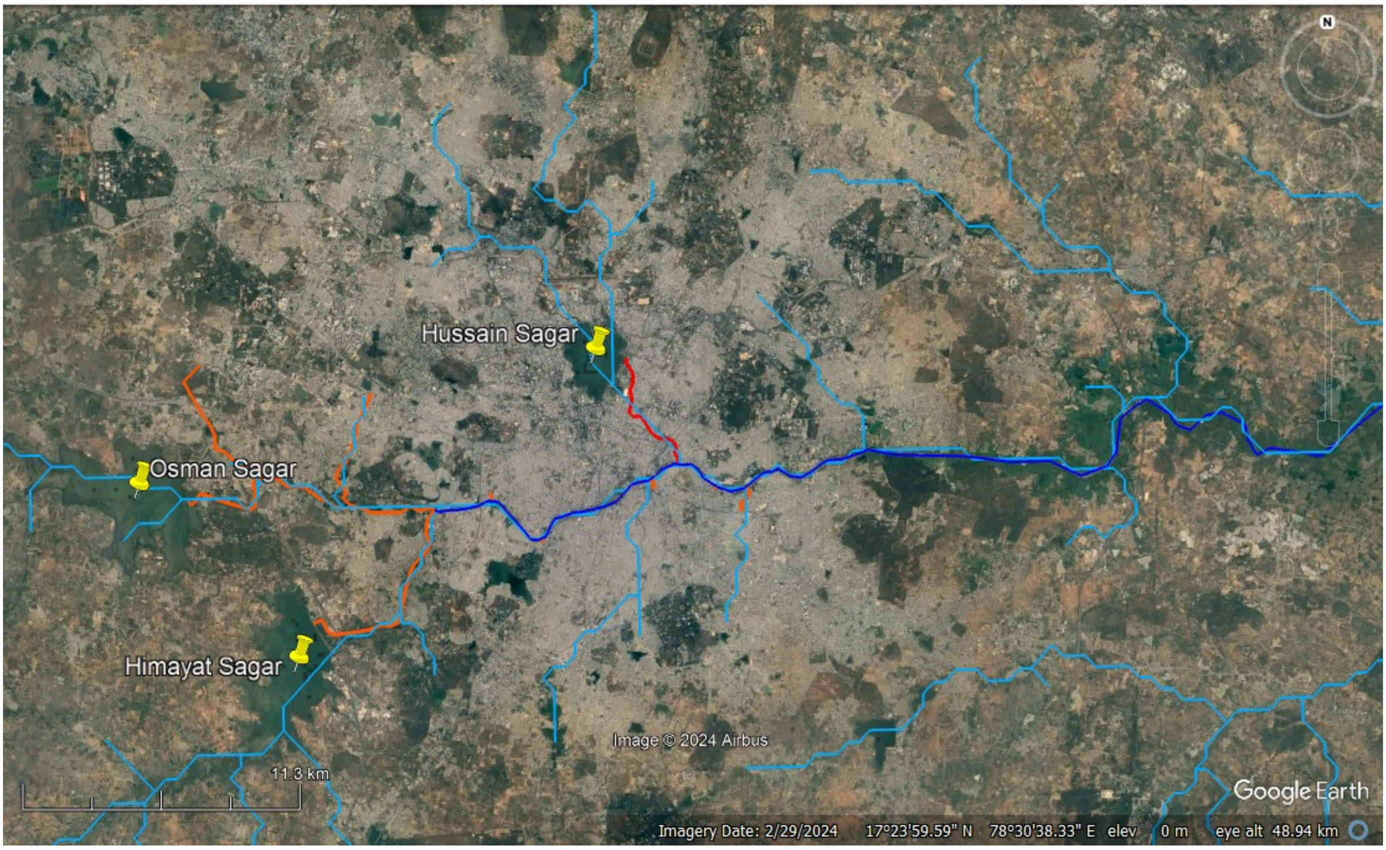


**Fig. S13.** Overlayed drainage network map from manual tracing and DEM delineation.

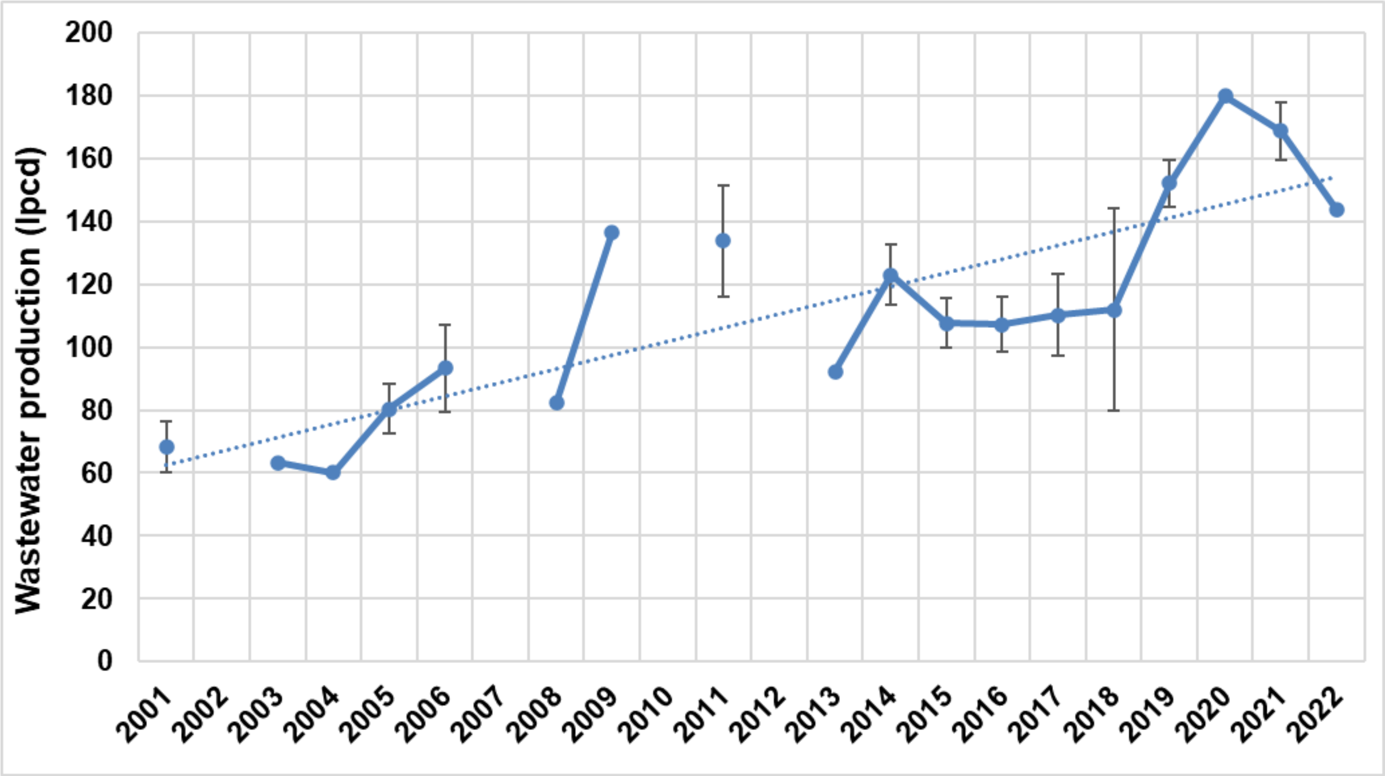


**Fig. S14.** The trend of estimated wastewater production (direct and indirect estimation methods combined) in litres per person per day (lpdc) in Hyderabad was roughly linear but showed substantial scatter. Data points were averages with standard error if more than one value was available in the given year.

**Table S4.** Physicochemical variables analysed in water samples.

| S. No. | Parameter | Unit | Method (APHA, 2017)[6] |
|---|---|---|---|
| 1 | Temperature | °C | |
| 2 | pH | | Potentiometry |
| 3 | Electrical Conductivity (EC) | µS/cm | Potentiometry |
| 4 | Total Dissolved Solids (TDS) | mg/L | Potentiometry |
| 5 | Dissolved Oxygen (DO) | mg/L | Polarographic probe |
| 6 | Turbidity (TUR) | NTU | Nephelometry (2130-B) |
| 7 | Total Suspended Solids | mg/L | Gravimetry (2540 D) |
| 8 | Volatile Solids | mg/L | Gravimetry (2540 E) |
| 9 | Chemical Oxygen Demand ($COD_{Total}$) (particulate and soluble) | mg/L | Titrimetry (5220-C) |
| 10 | Total organic carbon (TOC) | mg/L | TOC analyser |
| 11 | Total nitrogen (TN) | mg/L | TOC analyser |
| 12 | Ammonium ($NH_4^+ - N$) | mg/L | Ion Chromatography (Thomas et al., 2002)[7] |
| 13 | Nitrite ($NO_2^- - N$) | mg/L | Ion Chromatography (Thomas et al., 2002)[7] |
| 14 | Nitrate ($NO_3^- - N$) | mg/L | Ion Chromatography (Thomas et al., 2002)[7] |
| 15 | Total Phosphorus (TP) | mg/L | Vanadomolybdophosphoric Acid (4500-P C) |

**Table S5.** Physicochemical variables analysed in sediment samples.

| S. No. | Parameter | Unit | Method | Reference |
|---|---|---|---|---|
| 1 | pH | – | Potentiometry | (IS 2720 (Part26), 1987)[8] |
| 2 | Electrical Conductivity (EC) | µS/cm | Potentiometry | (IS 14767:2000, 2000)[9] |
| 3 | Moisture content (MC) | % | Gravimetry | (García-Ruiz et al., 1998)[10] |
| 4 | Organic Matter (OM) | mg/kg | | (Svendsen et al., 1993)[11] |
| 5 | Nitrate ($NO_3^- - N$ ) | mg/kg | Ion Chromatography | (Thomas et al., 2002)[7] |
| 6 | Ammonia ($NH_4^+ - N$ ) | mg/kg | Ion Chromatography | (Thomas et al., 2002)[7] |
| 7 | Total Organic Carbon (TOC) | mg/kg | TOC analyser | (Avramidis and Bekiari, 2021)[12] |
| 8 | Total Nitrogen (TN) | mg/kg | TOC analyser | (Avramidis and Bekiari, 2021)[12] |
| 9 | Total Phosphorus (TP) | mg/kg | Vanadomolybdophosphoric Acid (4500-P C) | (APHA, 2017)[6] |

**Table S6.** Oligonucleotides used for ARG and MGE detection by qPCR reactions.

| Target gene | Reference | Probe name | Oligonucleotide sequence 5'-3' | Conc. in reaction (nmol $L^{-1}$) | Amplicon size (bp) | Ann. $T_a$ (ºC) | Standard curve |
|---|---|---|---|---|---|---|---|
| **16S rRNA** | (Muyzer et al., 1993)[13] | q_338F | ACTCCTACGGGAGGCAGCAG | 200 | 198 | 60 | $R^2$= 0.996<br>Slope = -3.34<br>Efficiency (%) = 99.40 |
| | | q_518R | ATTACCGCGGCTGCTGG | | | | |
| ***qnrS*** | (Marti and Balcázar, 2013)[14] | q_qnrSrtF11 | GACGTGCTAACTTGCGTGAT | 200 | 117 | 60 | $R^2$= 0.998<br>Slope = -3.237<br>Efficiency (%) = 103.664 |
| | | q_qnrSrtR11 | TGGCATTGTTGGAAACTTG | | | | |
| ***uidA*** | (Frahm and Obst, 2003; Silkie et al., 2008)[15,16] | q_uidA-F | CGGAAGCAACGCGTAAACTC | 200 | 90 | 62 | $R^2$ = 0.999<br>Slope = -3.51<br>Efficiency (%) = 92.64 |
| | | q_uidA-R | TGAGCGTCGCAGAACATTACA | | | | |
| ***aph(3")-Ib*** | (Liang et al., 2021)[17] | q_aph(3")-Ib-F | ACTGGCAGGAGGAACAGGAGGGTG | 200 | 240 | 58 | $R^2$ = 1.000<br>Slope = -3.398<br>Efficiency (%) = 96.912 |
| | | q_aph(3")-Ib-R | CGTCCGGTAAGAAGTCGGGATTGA | | | | |
| ***sul2*** | (Pei et al., 2006)[18] | q_sul2-F | TCCGGTGGAGGCCGGTATCTGG | 200 | 192 | 62 | $R^2$ = 0.999<br>Slope = -3.51<br>Efficiency (%) = 92.64 |
| | | q_sul2-R | CGGGAATGCCATCTGCCTTGAG | | | | |
| ***bla<sub>NDM</sub>*** | Resistomap AY152 | q_blaNDM-F | GGCCACACCAGTGACAATATCA | 200 | 66 | 59 | $R^2$ = 0.997<br>Slope = -3.26<br>Efficiency (%) = 102.72 |
| | | q_blaNDM-R | CAGGCAGCCACCAAAAGC | | | | |
| ***bla<sub>CTX-M</sub>*** | (Marti et al., 2013)[19] | q_CTXM-F | CTATGGCACCACCAACGATA | 200 | 104 | 57 | $R^2$ = 0.999<br>Slope = -3.51<br>Efficiency (%) = 90.79 |
| | | q_CTXM-R | ACGGCTTTCTGCCTTAGGTT | | | | |
| ***intI1*** | (Barraud et al., 2010)[20] | q_intI-F | GATCGGTCGAATGCGTGT | 200 | 196 | 61 | $R^2$ = 0.999<br>Slope= -3.344<br>Efficiency (%) = 99.093 |
| | | q_intI-R | GCCTTGATGTTACCCGAGAG | | | | |
| ***ermF*** | Resistomap AY535 | q_ermF -F | TCTGATGCCCGAAATGTTCAAG | 300 | 170 | 57 | $R^2$ = 0.998<br>Slope = -3.227<br>Efficiency (%) = 104.113 |
| | | q_ermF-R | TGAAGGACAATTGAACCTCCCA | | | | |
| ***tetW*** | (Walsh et al., 2011)[21] | q_tetW -F | CGGCAGCGCAAAGAGAAC | 200 | 59 | 57 | $R^2$ = 0.986<br>Slope = -3.28<br>Efficiency (%) = 101.94 |
| | | q_tetW -R | CGGGTCAGTATCCGCAAGTT | | | | |
